# Insights into spinach domestication from genomes sequences of two wild spinach progenitors, *Spinacia turkestanica* and *S. tetrandra*

**DOI:** 10.1101/2023.11.08.566323

**Authors:** Hongbing She, Zhiyuan Liu, Zhaosheng Xu, Helong Zhang, Jian Wu, Xiaowu Wang, Feng Cheng, Deborah Charlesworth, Wei Qian

**Author notes:** These authors contributed equally to this work (Hongbing She, Zhiyuan Liu). Correspondence: Wei Qian, Deborah Charlesworth, Feng Cheng.

## Abstract

Cultivated spinach (*Spinacia oleracea*) is a dioecious species (with male and female flowers on separate individuals). We report high-quality genome assemblies for its two closest wild relatives, *S. turkestanica* and *S. tetrandra*, which are also dioecious, to study the genetics of spinach domestication. Using a combination of genomic approaches, we assembled genome sequences of both these species, and analysed them in comparison to the previously assembled *S. oleracea* genome. These species diverged approximately 6.3 million years ago (Mya), while cultivated spinach split from *S. turkestanica* (its probable direct progenitor) 0.8 Mya. A common feature of all three species is that all six chromosomes include very large gene-poor, repeat-rich regions. In *S. oleracea*, these correspond with pericentromeric regions with very low recombination rates in both male and female genetic maps, and we infer that the similar pericentromeric regions in the wild species also recombine rarely. Although these regions include a low proportion of *Spinacia* genes, many genes are nevertheless within them, and they must be considered when analyzing selection during domestication. As a first approach to the difficult question of detecting genes involved in spinach domestication, we characterized 282 structural variants (SVs) whose frequencies are higher in a set of spinach accessions than in the wild species, suggesting that they mark genome regions that have been selected during domestication. These regions include genes associated with leaf margin type and flowering time. We also describe evidence that the downy mildew resistance loci of cultivated spinach are derived from introgression from both wild spinach species.

## Main text

It is estimated that between 1,000 and 2,500 plant species have been semi- or fully domesticated (Purugganan, 2019), and that today humans rely on many few crops that were domesticated more than 10,000 years ago (Ross-Ibarra et al., 2007). Although many advanced technologies are applied for agronomic improvements (Hickey et al., 2019), low genetic diversity often due to genetic bottlenecks at initial domestication often hinders these efforts (Mascher et al., 2021), and wild relatives can be a valuable sources of genetic factors for traits desired in cultivated crops (Lin et al., 2014; Tian et al., 2019). Genome sequences and assemblies of crop wild relatives have therefore been increasingly used to explore adaptation to different environmental conditions, domestication, and traits of interest for crop breeders (Sun et al., 2020; Zhang et al., 2023).

Bottlenecks during domestication severely hinder efforts to infer the genetics of crop improvement. Furthermore, bottlenecks lead to associations between variants (linkage disequilibrium, or LD) across larger recombination distances than expected in populations with constant size, hindering detection of selection based on unusual LD patterns, such as Hamblin et al., (Hamblin et al., 2006). Importantly, genetic mapping in cultivated spinach has shown that each chromosome includes a very large region in which recombination rates are very low. These regions extend over physical distances around the centromeres of chromosomes, and are termed “pericentromeric regions”.

To test whether pericentromeric regions in plants and animals undergo rare recombination, or are recombinationally completely inactive, population genomic studies are needed, including analyses of the decay of LD (Chen et al., 2022; McVean et al., 2002). In non-recombining or rarely recombining regions, LD can extend over large physical distances, greatly hampering analyses intended to detect selected loci: most loci across the region will have nearly identical evolutionary histories, and a change in the frequency of a favoured allele at one locus will change the frequencies of alleles of many other genes all across its linked region (Nielsen et al., 2005; Stephan, 2019). The problems are compounded by the tendency for such rarely recombining regions to have high repetitive sequence densities, including transposable elements (TEs) and satellites (Charlesworth et al., 1994; Dolgin and Charlesworth, 2008), which can make reliable genome assembly extremely difficult.

The cultivated spinach has two close wild relatives, *S. turkestanica* Iljin and *S. tetrandra* Stev. (Ribera et al., 2020). Phylogenic inferences using nuclear and chloroplast genome sequences demonstrated that *S. turkestanica* is closer to *S. oleracea* than is *S. tetrandra* (She et al., 2022; Xu et al., 2017). Both wild relatives possess genetic resources of value for improvement of cultivated spinach, including resistance to pests and disease (Treuren et al., 2020). Here, we used differences in genome organization (structural variants) between wild and cultivated spinach to detect genome regions involved in spinach domestication.

We combined a multifaceted approach for chromosome-scale assembly of *S. turkestanica* and *S. tetrandra* genomes (Tables S1 and S2). We *de novo* assembled a 963 Mb genome for a male of the closer outgroup species, *S. turkestanica* (with contig N50=3.49 Mb) using 56 Gb of Oxford Nanopore sequencing (ONT) reads and 56 Gb of Illunima reads. The assembly size is close to the estimated genome size of 998 Mb using k-mer distribution analysis (Fig. S1a). In this assembly, named Tu17S31XY, contigs totalling 931 Mb (96.60%) were anchored to the correct number of chromosomes, six, based on a reference-guided approach using the cultivated spinach male reference assembly, Sp_YY_v1 (Fig. 1, Tables S1 and S2).

**Figure 1.**
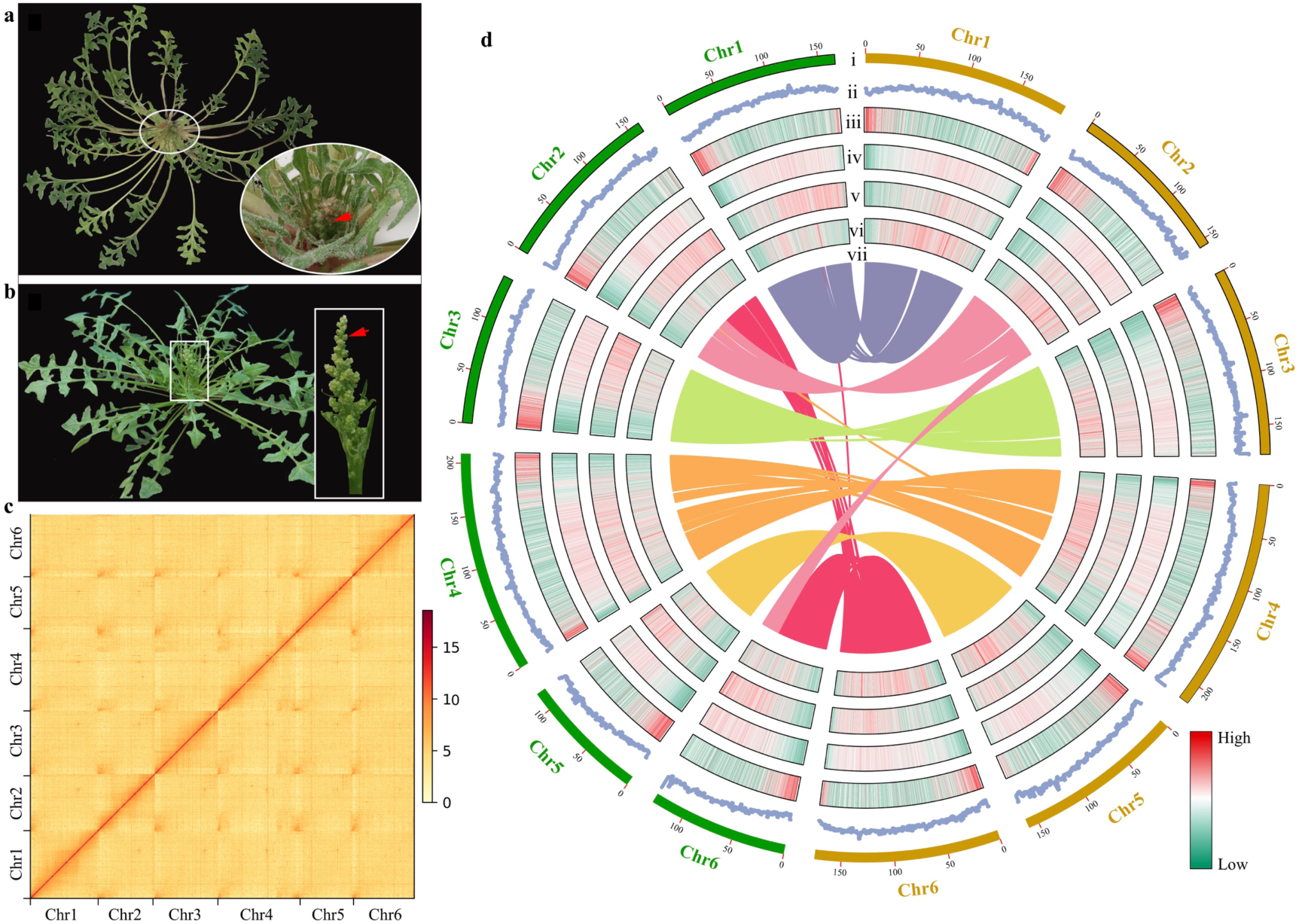
Morphology of individuals and inflorescences of the two wild spinach species, illustrating the differences between them, and the species’ genome features. Morphology of (a) *Spinacia tetrandra* and (b) *S. turkestanica*. Red arrows represent stamens. **(c)** Hi-C contact heatmap of the *S. tetrandra* assembly at 500-kb resolution. **(d)** The genomic landscape of the two assemblies. (i) the six pseudochromosomes of *S. turkestanica* (green bars) and *S. tetrandra* (brown bars) assemblies, (ii) the guanine-cytosine (GC) contents, (iii) to (vi) gene densities, densities of the *Copia*, and *Gypsy* element transposable elements (TEs), (vii) syntenic blocks in the two wild spinach genomes. The window size for tracks (ii) to (vi) was 100 kb.

For the less closely related wild species, *S. tetrandra*, a raw assembly of 1.45 Gb was achieved using 58 Gb ONT reads and 50 Gb Illumina reads. A high proportion of sites in our reads, 1.77%, were heterozygous (Fig. S1b), so we removed redundant sequences using Purge_haplotigs (Roach et al., 2018); the resulting monoploid assembly (an assembly with parental alleles randomly switching in a contig) size was 1.14 Gb, with a contig N50 of 5.38 Mb. After Hi-C scaffolding, 94.50% of the *S. tetrandra* contigs were again anchored to six chromosomes in an assembly named Te17S22XY (Fig. 1c, Table 1, Tables S1–S3).

**Table 1.** Statistics of the genomic assembly and annotation for the two wild spinach species.

| Items | *S. turkestanica* | *S. tetrandra* |
| --- | --- | --- |
|  | Tu17S31XY | Te17S22XY |
| Number of contigs | 805 | 431 |
| Contig N50 (Mb) | 3.49 | 5.38 |
| Assembly size (Mb) | 963 | 1,143 |
| Anchored chromosome (Mb) | 931 | 1,081 |
| LTR assembly index (LAI) | 18.08 | 18.12 |
| Repetitive sequences (%) | 74.16 | 75.55 |
| Number of annotated genes | 26,631 | 29,521 |
| Mean transcript length (bp) | 4787 | 5317 |
| Mean exon length (bp) | 258 | 257 |
| Mean exon per mRNA | 5.20 | 5.06 |
| BUSCO completeness of annotation (%) | 97.8 | 98.6 |

Benchmarking Universal Single-Copy Orthologs (BUSCO (Waterhouse et al., 2017)) evaluations of both wild spinach assemblies, using embryophyta_odb10 database, yielded estimates of 97.8% and 98.6% completeness for the *S. turkestanica* and *S. tetrandra* assemblies, respectively (Table S3). The accuracy and completeness of the assemblies were further supported by high values (18.08 and 18.12) of the Long terminal repeat (LTR) LTR assembly index (LAI, (Ou and Jiang, 2018)) and high mapping rates of ∼50× Illunima short reads (≥ 99.65%), RNA-seq (96.30%), and Iso-seq (99.80%) from five tissues. (Table 1, Tables S4, S5, and Fig. S2).

We annotated 26,631 and 29,521 protein-coding genes in the *S. turkestanica* and *S. tetrandra* assemblies, respectively (Table 1 and Table S6). More than 96% of these genes could be annotated (Table S7). We also identified 6,618 and 7,741 non-coding RAN containing ribosomal RNAs (rNRAs), transfer RNA (tRNAs), microRNAs (miRNAs), and small nuclear RNAs (snRNAs) of *S. turkestanica* and *S. tetrandra* assemblies, respectively (Table S8).

To check the previously inferred evolutionary history of the three *Spinacia* species, we identified 23,568 gene families were identified across the seven species (Fig. 2a, b, Table S9). Fig. 2c shows a time-calibrated phylogenetic tree using the 812 single-copy genes that are shared among all seven species (Table S10). The divergence times between the three *Spinacia* species, indicate that *S. turkestanica* and *S. oleracea* differentiated from *S. tetrandra* approximately 6.3 million years ago (Mya), and that *S. oleracea* and *S. turkestanica* diverged much more recently (∼0.8 Mya, see Fig. 2c), consistent with *S. turkestanica* being the ancestor of cultivated spinach, as previously inferred using chloroplast genome sequences (Ma et al., 2022; She et al., 2022).

**Figure 2.**
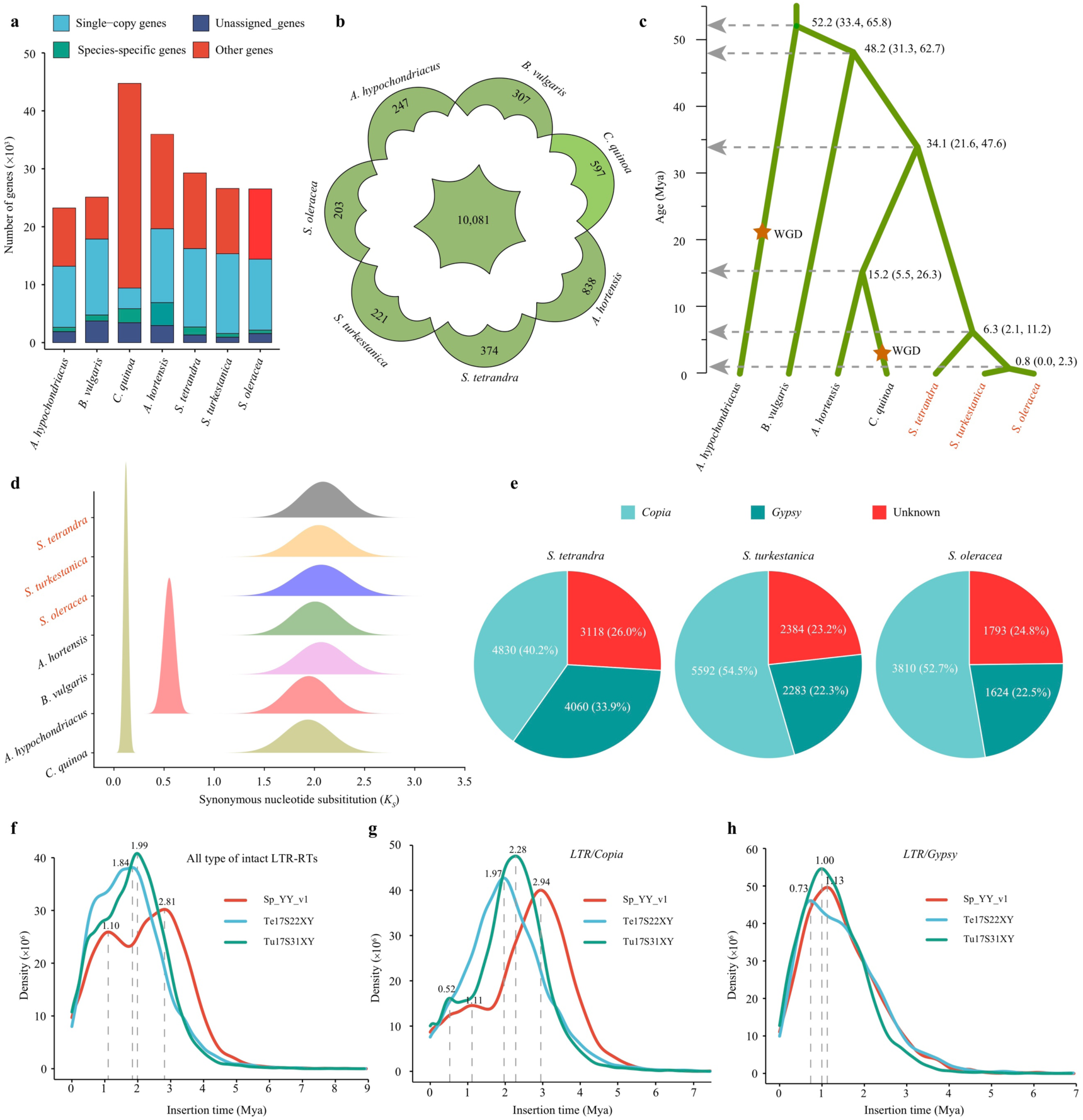
Gene family numbers, relationships, divergence time estimates, and intact-LTR-RT insertion time analyses. **(a)** The gene numbers in each category in three *Spinacia* species and four representative outgroup species, quinoa (*Chenopodium quinoa*), garden orache (*Atriplex hortensis*), sugar beet (*Beta vulgaris*), and amaranth (*Amaranthus hypochondriacus*). **(b)** Shared and species-specific gene families in the same species. **(c)** Phylogenetic relationships and estimated divergence times based on 812 single-copy genes detected in all seven species (see the text); the 95% highest posterior density ranges (in brackets) are shown at each node. **(d)** Distribution of synonymous divergence (*K_S_*) values between paralogs that are syntenic in the three *Spinacia* species and the outgroup species. **(e)** The numbers of intact LTR-RTs in the three *Spinacia* specie genomes. **(f–h)** Distribution of intact LTR-RT element insertion times. Insertion times were inferred assuming a mutation rate of 7.0 × 10^−9^ substitutions per site per year as described in Xu et al. (2017). Mya: million years ago.

We nevertheless tested for possible WGD events by estimating synonymous nucleotide divergence (*Ks*) values between paralogous genes, using the WGDI software (Sun et al., 2022). The *Ks* distributions in all seven species had only single peaks (Fig. 2d), reflecting an ancient γ whole-genome triplication (WGT) event shared by the three *Spinacia* species and the four outgroup species, and by most eudicotyledonous species so far studied (Jaillon et al., 2007). We conclude that the three *Spinacia* species are not recent polyploids, and that, after the WGT event, most genes have become diploid again, by loss of copies.

Genome-wide repeat densities are similar in all three species, ranging between 74 and 75% (Table 1) (She et al., 2023). Both the wild spinach species have high transposable element (TE) densities, with long terminal repeat elements (LTRs) forming 42.71% and 44.39% of the two genomes (Table 1 and Table S11). These densities are similar to those found in *S. oleracea* (She et al., 2023; Xu et al., 2017). Intact LTR-RTs in the three *Spinacia* species yielded similar results (Fig. 2e). However, full length *Copia* and *Gypsy* have similar densities in *S. tetrandra*, whereas *Copia* densities are almost twice those of *Gypsy* elements in the closer relatives *S. turkestanica* and *S. oleracea* (Table S11). Insertion times, estimated using divergence between the LTR sequences, revealed *LTR/Copia* burst event in *S. turkestanica* (0.52 Mya) and *S. oleracea* (1.11 Mya), which might account for their larger full-length *Copia* than *Gpysy* content (Fig. 2f–h, Fig. S3).

We previously found repeat-rich regions at one end of each *S. oleracea* autosome, and in the middle region of the sex chromosome pair (She et al., 2023). Table 2 shows that all three species, including the two wild spinach relatives, have similar extensive gene-poor/TE-rich regions (Fig. 1). These are probably pericentromeric regions (Table 2) with rare recombination, which allows the accumulation of repetitive sequences (Fig. S4). The gene densities indeed change sharply between the pericentromeric (mean 17 genes per Mb) and non-pericentromeric regions (mean 53 genes per Mb) among the three *Spinacia* species (Table 2).

**Table 2.**
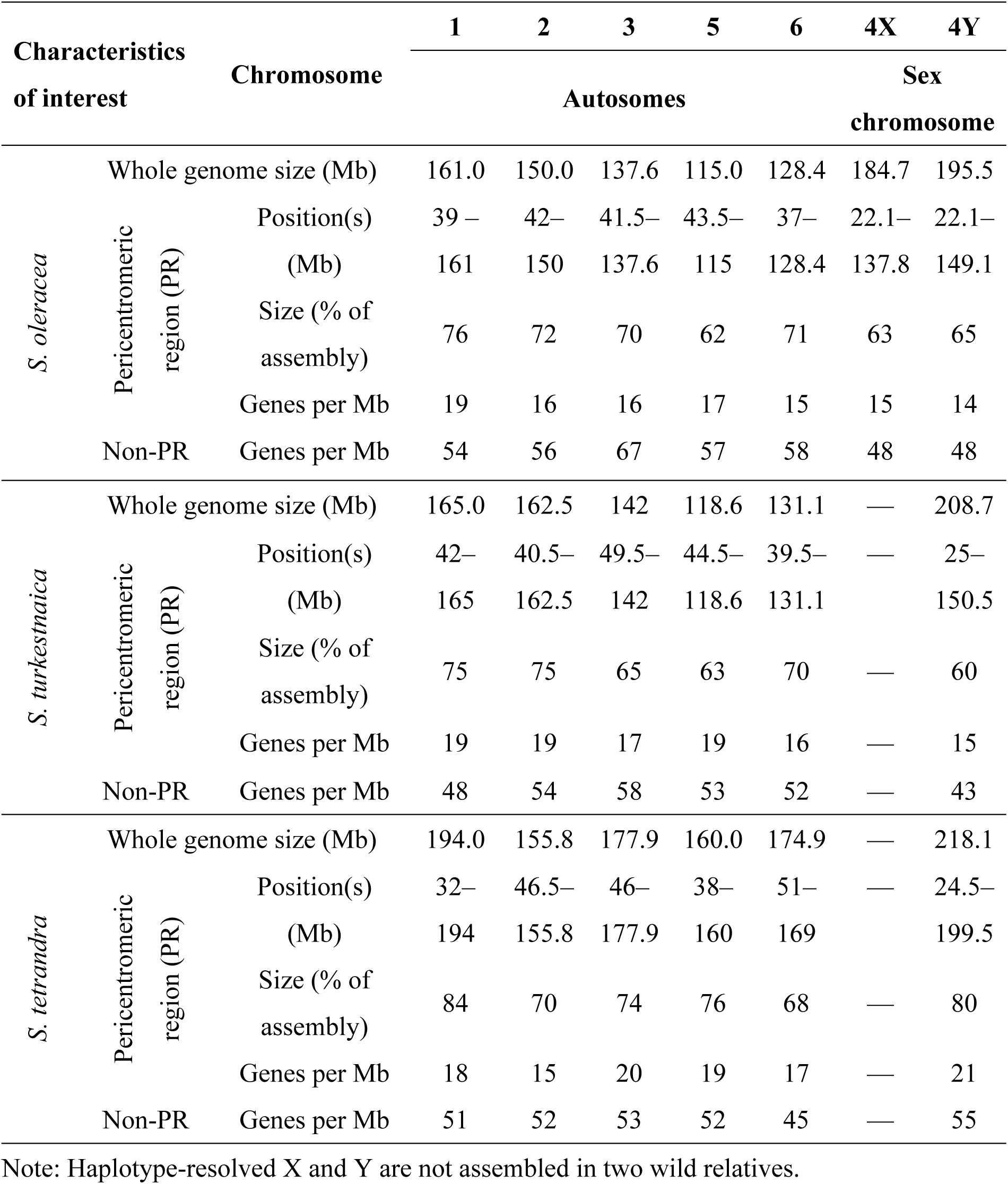
Characteristics of pericentromeric and non-pericentromeric regions among the three *Spinacia* species.

Regions with low recombination rates often have low nucleotide diversity. We estimated genome-wide nucleotide diversity (expressed as π values for all site types), and compared the values between pericentromeric and non-pericentromeric regions using 138,697 high-quality SNPs (< 0.05 missing data, MAF > 0.05) using sequences from 13 *S. tetrandra* accessions mapped to the Te17S22XY assembly. For every chromosome, the mean π value for pericentromeric regions is significantly lower than for non-pericentromeric regions (Fig. 3a, Wilcoxon tests, *P* < 0.01, Table S12). The mean π across all the pericentromeric regions is 0.05×10^-3^, versus 0.13×10^-3^ for non-pericentromeric regions of all chromosomes.

**Figure 3.**
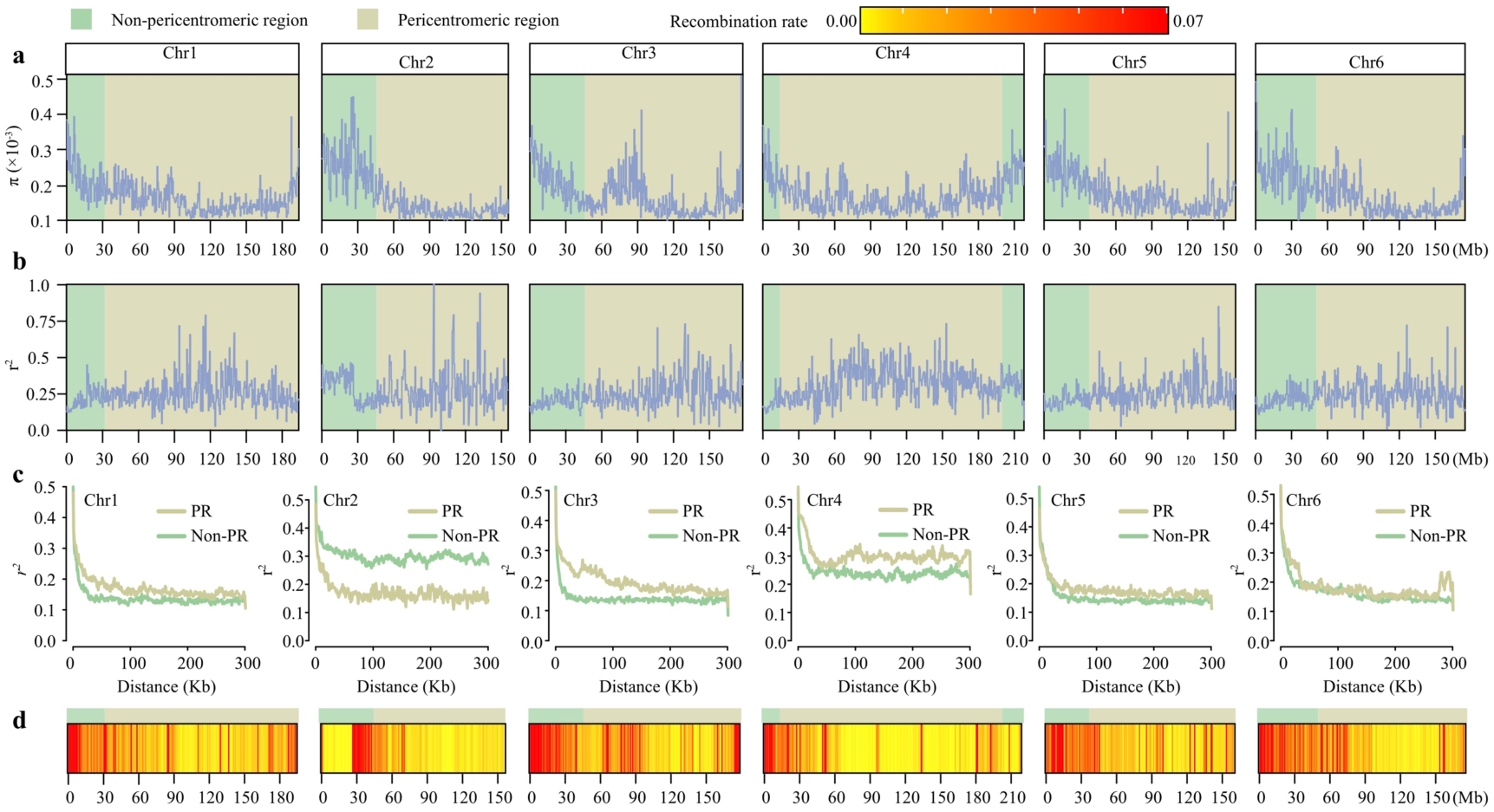
Distribution of nucleotide diversity (π), recombination rate and linkage disequilibrium (LD) values for genes on each chromosome in 13 *S. tetrandra* accessions, with their positions in the Te17S22XY assembly shown on the x axis. The green and yellow regions are non-pericentromeric and pericentromeric, respectively. π **(a)** and LD **(b)** (measured as pairwise *r^2^* values within 50-kb windows) were estimated in 500-kb sliding windows. **(c)** LD decay, showing the generally faster decay in the non-pericentromeric (Non-PR) than the pericentromeric regions (PR). **(d)** Heat map of recombination rate on each chromosome (red indicates higher and yellow indicates lower rates).

If recombination is indeed rare, the pericentromeric regions should also show high linkage disequilibrium (LD). We measured LD as *r^2^* values, which are indeed much higher in the pericentromeric regions than other genome regions (despite considerable variation, especially within the pericentromeric regions, Wilcoxon tests with *P* < 0.01, see Fig. 3b). Pericentromeric regions also show slower LD decay than with non-pericentromeric ones (Fig. 3c). The *r*^2^ value to which LD decays depends on the number of gametes sampled (for autosomes, this is double the number of individuals analyzed), and the very similar levels at large distances (300 kb in Fig. 3c), for most chromosomes, and for both regions, as therefore expected. The differences for the sex chromosome, chr4, reflects their smaller number of gametes sampled, together with the fact that crossovers do not occur across the pericentromeric region, which is Y-linked in *S. tetrandra* males, as well as being rare or absent in females. Chromosome 2, however, is an exception, as part of the non-pericentromeric region shows high LD and slow LD decay (Fig. 3b, c, Table S13).

The observation that LD levels between SNPs at large physical distances for the pericentromeric regions are similar to those for the highly recombining regions (Fig. 3c) suggests that recombination occurs in the pericentromeric regions, albeit rarely. This is supported by quantitative estimates using LDHelmet (Chan et al., 2012), which consistently detect recombination within the pericentromeric regions (Fig. 3d). The high LD in these regions is therefore not due to a complete absence of recombination, other than the region of about 25 Mb of the left-hand end of chr2 mentioned above, which shows a complete absence of recombination, perhaps reflecting an inversion polymorphism similar to the one detected in barley (Jayakodi et al., 2020). If recombination is suppressed in heterozygotes for different arrangements, an inversion segregating at a high frequency within this species could explain this result.

The diversity differences between the pericentromeric and recombinationally active genome regions might reflect mutation rate differences, since higher recombination rates tend to be correlated with increased mutation rates (Bussell et al., 2006; Filatov and Gerrard, 2003; Hellmann et al., 2003). We tested this by estimating divergence between *S. tetrandra* and the other *Spinacia* species. Supplementary Fig. 5 shows that high diversity regions also show high divergence, suggesting that mutation rates differ in different genome regions. To distinguish between different possible causes, we analysed GC content, as high recombination rates tend to be correlated with increased GC content, due to GC-biased gene conversion (Marais and Galtier, 2003). In *S. tetrandra*, the synonymous site diversity differences described above suggest that the recombinationally active regions are physically large. Their recombination rates per megabase might therefore not be expected to be exceptionally high, and so the GC content should not differ much between the different regions. Indeed the GC content of introns is quite uniform, with no pronounced difference between the regions (Fig. S5). We therefore conclude that, although the higher nucleotide diversity values in the non-pericentromeric regions are probably caused by higher mutation rates in those regions, not due to high GC content (Arbel-Eden and Simchen, 2019). The mechanisms causing elevated mutagenicity in meiosis are currently unknown (Arbel-Eden and Simchen, 2019).

To test whether diversity differences are consequences of recombination rate differences other than via mutation rates, one can correct for this effect by scaling synonymous site diversity estimates for genes by their synonymous site divergence estimates (Begun and Aquadro, 1992). The raw and corrected diversity value of single-copy gene in 13 *S. tetrandra* accessions shared similar distribution patterns, suggesting that mutation rates did not affect diversity (Fig. S6, Table S14). Diversity in recombinationally inactive pericentromeric regions can also be reduced due to selective sweeps when advantageous mutations in such regions spread, and also by selection eliminating deleterious mutations, which reduces the region’s effective population size (Charlesworth et al., 1993).

The *S. tetrandra* assembly is not collinear with the *S. turkestanica* and *S. oleracea* assemblies, which are highly collinear with each other (Fig. 4a). Sequence alignment between the *S. turkestanica* and *S. tetrandra* assemblies suggests that the sex chromosome, chromosome 4, underwent more inversions > 1Mb than the autosomes (the mean inversion numbers per 10 Mb are 1.33 and 0.29, respectively) (Fig. 4a, Fig. S7, Table S15). Finally, Hi-C analysis detected a large reciprocal translocation between the two wild spinach assemblies, involving the repeat-rich terminal regions (a 21.52 Mb chromosome 2 region in the *S. tetrandra* assembly, and a 53.21 Mb chromosome 6 region) (Fig. 4 b, Table S16).

**Figure 4.**
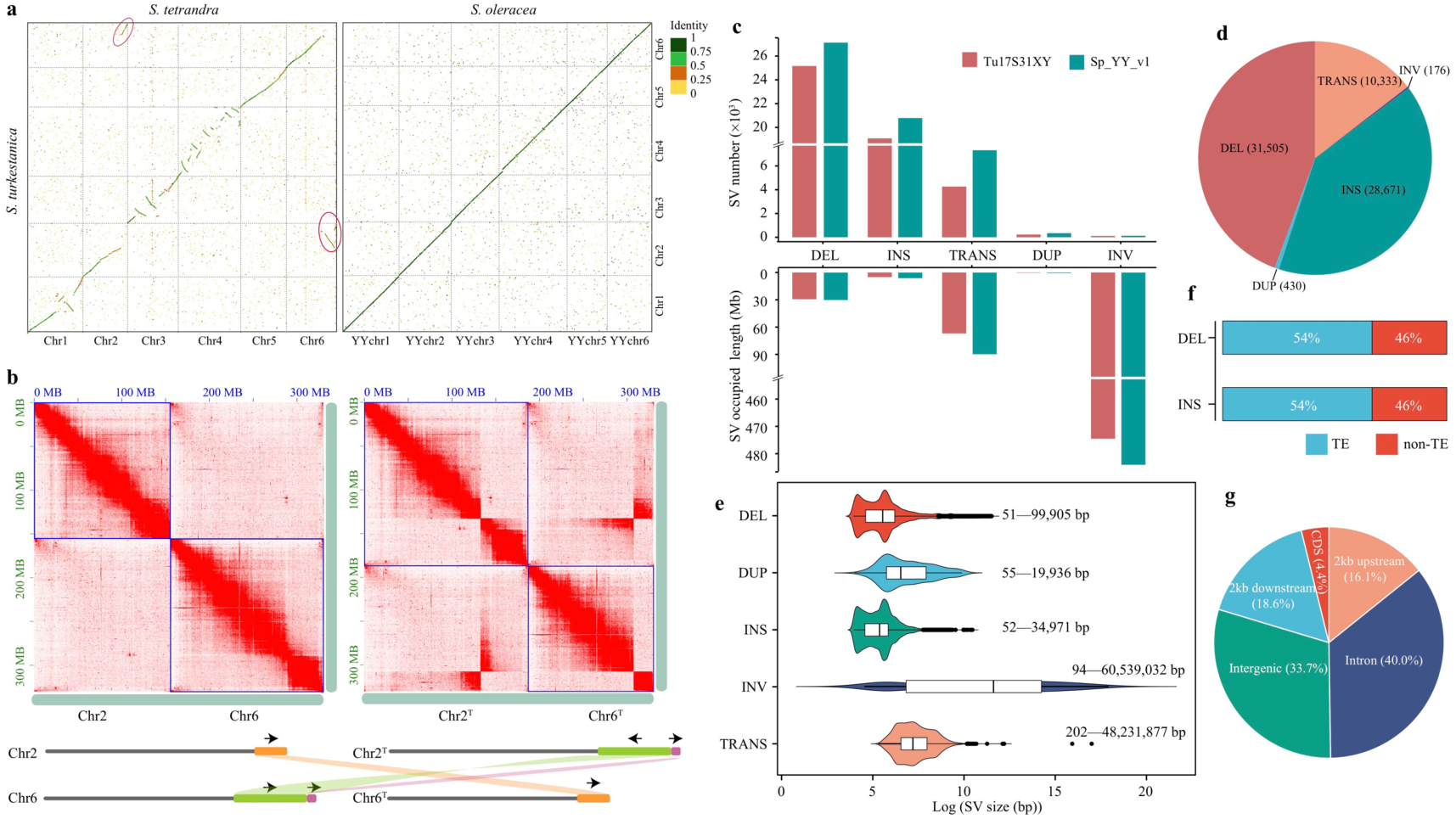
Inference and characterization of spinach SVs. **(a)** Dot-plot of Tu17S31XY (*S. turkestanica*) and Te17S22XY (*S. tetrandra*), Sp_YY_v1 (*S. oleracea*) assemblies. The two red symbols outline regions where a reciprocal translocation is detected between the *S. tetrandra* and *S. turkestanica* genomes involving chromosomes 2 and 6. **(b)** Hi-C interaction heatmaps of chromosomes 2 and 6 at 500-kb resolution. Chr2 and Chr6 denote chromosomes from the *S. tetrandra* Te17S22XY assembly, while Chr2^T^ and Chr6^T^ indicate the translocated Chr2 and Chr6 chromosomes (as diagrammed at the bottom of part **b**). **(c)** The numbers of five types SVs in the Tu17S31XY and Sp_YY_v1 assemblies. **(d)** The percentages of non-redundant SVs divided into the five types described in the text between the Te17S22XY and Tu17S31XY/Sp_YY_v1 assemblies. **(e)** Size distributions of the five SV types. **(f)** The percentage of non-redundant insertions and deletions overlapping TEs. **(g)** Percentages of non-redundant SVs overlapping different genomic regions.

We compared the *S. oleracea* and *S. turkestanica* reference genome sequences (Sp_YY_v1 and Tu17S31XY, respectively) against our new *S. tetrandra* reference (Te17S22XY), representing the likely ancestral state). We used both read-mapping-based and assembly-based approaches to identify SVs > 50 bp, including deletions, insertions, duplications, inversions, and translocations additional to the reciprocal one just described. Among the five SV types, deletions are the most frequent, while inversions were the rarest SV type, but occupied the largest amounts of sequence, 483.7 Mb and 475.0 Mb in the *S. oleracea* and *S. turkestanica* assemblies, respectively (Fig. 4c), or approximately half of these species’ total genome sizes; 54 non-redundant inversions ≥ 1 Mb, especially on chromosomes 1, 3 and 4, correspond to a total of 522.9 Mb (Table S15, Fig. S8). Compared with the Sp_YY_v1 assembly, the outgroup *S. tetrandra* consistently differs more in terms of SVs (across different SV types, with a total of 55,727 SVs, occupying a total of 614.3 Mb) than the Tu17S31XY one, with 48,849 SVs and 598.5 Mb (Fig. 4c, Table S17). The numbers of differences are consistent with the relationships of the two wild species with cultivated spinach previously estimated by phylogenetic analyses of sequences (Ma et al., 2022; She et al., 2022; Xu et al., 2017).

We then merged the SVs in the *S. turkestanica* and *S. oleracea* assemblies into a non-redundant set of 71,115 SVs that differ from the arrangement in the outgroup assembly. These SVs cover about 689.9 Mb in total, and include 31,505 deletions, 28,671 insertions, 176 inversions, 10,333 translocations, and 430 duplications (Fig. 4d, e, Table S18, Figs. S9 and S10. As expected, many of the insertion/deletion differences (54%) overlap with TEs (Fig. 4f, Fig. S11). We identified 309 SV-rich regions (Table S19), which were mainly found at the chromosome ends, rather than in the pericentromeric regions (Fig. S7). On the autosomes, which are acrocentric, slight enrichments were seen at most of the centromeric ends, but SVs were mainly concentrated at the non-pericentromeric (high gene density) end of each assembly (see Figure 1).

Only 4.4% of the SVs overlap coding sequences (CDS) (Fig. 4g), while 40.0% were found in introns. 13.3% of genes in the *S. tetrandra* assembly had at least one SV overlapping the 2 kb regions upstream of the start codon, or downstream of the stop codon. We term these genes “SV-related genes”. Across the pericentromeric regions of all chromosomes, 50% of genes were SV-related, versus 71% for non-pericentromeric regions. A total of 4,623 genes were found in these SV-rich regions. Notably, a 4.2-Mb region in the non-pericentromeric region of chromosome 1 (0–4.2 Mb) with 2,955 SVs, includes 30 resistance genes, eight of them being NBS family genes (Fig. S13, Table S21).

We genotyped 982,999 high-quality SNPs and 71,115 SVs ascertained in our long-read assemblies (as described above), in a set of 13 *S. tetrandra*, 22 *S. turkestanica* and 59 *S. oleracea* accessions from different sources (Table S22). A phylogenetic tree, and principal component analyses (PCA) based on the SNPs in all 94 *Spinacia* accessions is consistent with the relationships previously inferred (Fig. S14). A tree based on presence/absence of the SVs also consistently divided these *Spinacia* accessions into three groups corresponding with the three species (Fig. 5a, Fig. S15); the tree is almost star-shaped, and the two *S. turkestanica* accessions possible exceptions, Sp20 and Sp83, are placed among other individuals of this species.

**Figure 5.**
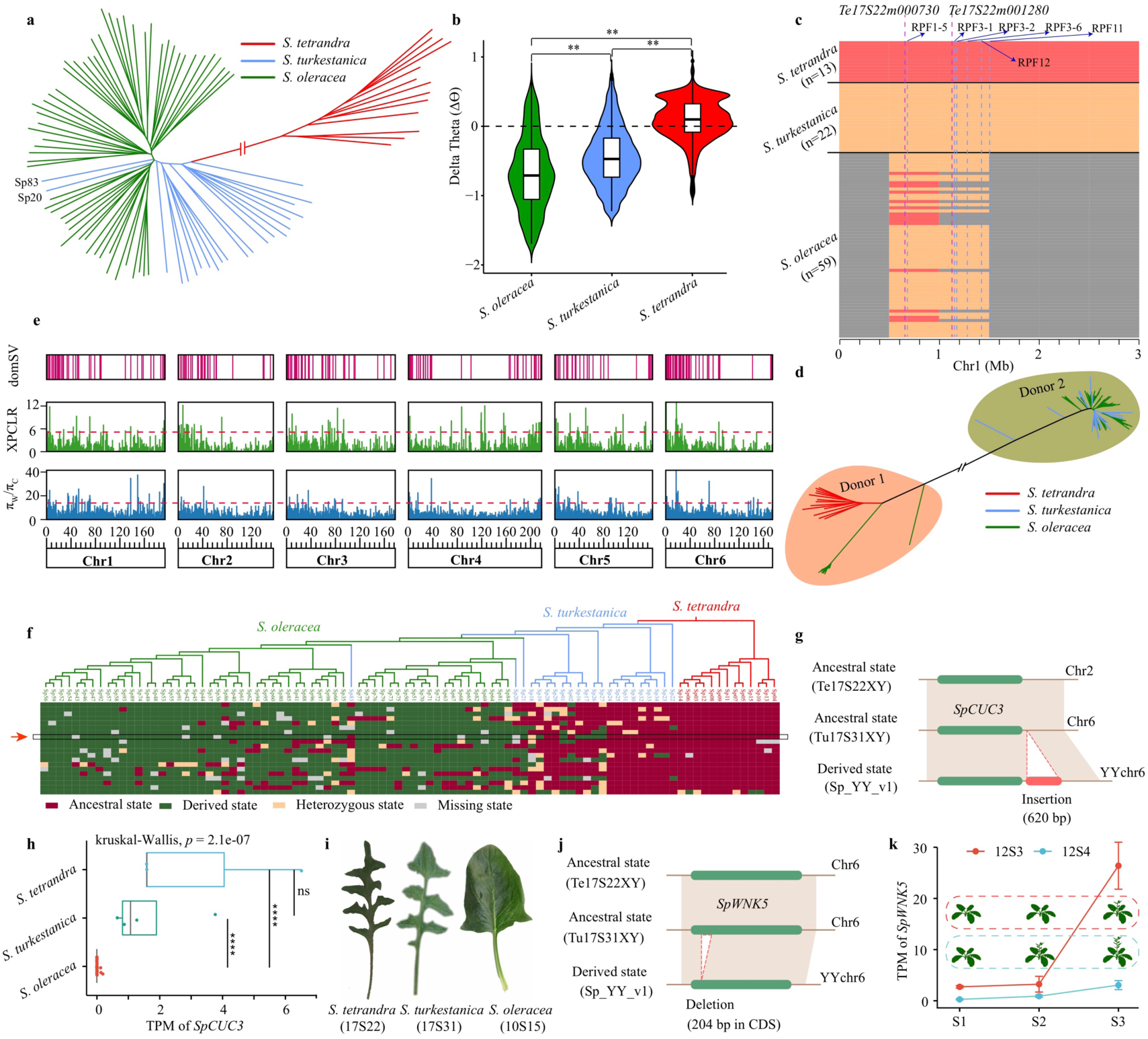
Phylogeny, introgression, and domestication in cultivated spinach and its wild relative species. Phylogenetic tree **(a)** of 13 *S. tetrandra*, 22 *S. turkestanica*, and 59 *S. oleracea* accessions based on SVs (Insertion and deletion). **(b)** Distributions of Delta Theta (Δϴ_s_) values for fourfold degenerate sites from 94 *Spinacia* accessions. **indicates significance (*P* < 0.01) by Kruskal–Wallis tests. **(c)** A representative introgression region that contains downy mildew (DM) resistance locus from two spinach wild progenitors (*S. tetrandra* and *S. turkestanica*) into the *S. oleracea*. Species-specific genome regions are shown in different colors. The dotted blue lines with blue reflect markers tightly linked to DM resistance genes, *RPF1*–*RPF3*, *RPF11*, and *RPF12*. Two candidate genes, *Te17S22m000730* and *Te17S22m1280*, of *RPF1* and *RPF2/3*, respectively, were colored with purple. **(d)** A neighbor-joining tree is constructed using SVs within the representative introgression region. **(e)** XPCLR, and nucleotide diversity ratio (π_W_/π_C_), are used for selection analysis in *S. oleracea.* Vertical dashed lines indicate genome-wide threshold of selection signals (XPCLR > 4.74 and π_W_/π_C_ > 13.73). Domestication SV (domSV) correspond to Fig. S22. **(f)** The genotype of top 20 selection signatures in the 94 *Spinacia* accessions. For an SV state that is the same as *S. tetrandra* is defined as the ancestral state, otherwise that is referred as derived state. **(g)** An insertion occurred at downstream of *SpCUC3* in *S. oleracea* (10S15) compared with two wild relatives. **(h)** The two wild relative accessions share significant higher expression level of *SpCUC3* compared with *S. oleracea* accessions. \*\*\*\**P* < 2.1×10^-7^, Kruskal-Wallis test. ns: no significant. **(i)** The different leaf type among the three *Spinacia* species. **(j)** A deletion occurred at coding region of SpWNK5 in *S. oleracea* (10S15) compared with two wild relatives. (k) The expression patters of the *SpWNK5* in the *S. oleracea*, 12S3 (late flowering) and 12S4 (early flowering). S1: neither plant is at flowering stage; S2: 12S4 is at flowering stage while 12S3 is not; S3: both plants are at flowering stage. The cartoon within the red and blue dotted ellipses mean three developmental stages for 12S3 and 12S4, respectively.

With sequence data, it is preferable to compare the two diversity measures, by computing Δϴ = (ϴ - π)/ϴ (or, equivalently 1 - (π/ϴ) (Becher et al., 2020; Langley et al., 2014; Tajima, 1989), which normalizes the differences between the two diversity measures by the total diversity, making its value less dependent on the total diversity for the sample (Jackson et al., 2017; Langley et al., 2014). We calculated Δϴ for fourfold degenerate sites, which should most closely approximate neutrality, and therefore be suitable for testing for deviations caused by recent population size changes; this is termed Δϴ_S_. The Δϴ_S_ values are much lower in *S. oleracea* (with mean -0.67 ± 0.53) than in the wild species (-0.44 ± 0.42 in *S. turkestanica*, and in *S. tetrandra* they are close to the neutral expectation (0.09 ± 0.31, see Fig. 5b), consistent with the previous conclusion that *S. oleracea* does not appear to have undergone a recent bottleneck.

The low Δϴ_S_ values in *S. oleracea*, might also reflect introgression of genomic regions from wild species, producing an excess of variants at intermediate or high frequencies, compared with the expectation at neutral equilibrium. We therefore tested for introgression using a likelihood ratio test described by Mcnally et al. (2009). This identified only six introgressed regions, most or all (as expected) from the closer relative, *S. turkestanica*, and occupying a total of 3.5 Mb (Fig. S16, Table S23). The same number of likely introgression regions was inferred by Ma et al. (2022) though their regions are larger. A total of 154 genes were detected within the regions detected here (Table S24).

Notably, one region was identified at 0.5–1.5 Mb of the *S. oleracea* chromosome 1 (Fig. 5c), overlapping with the downy mildew (DM) resistance loci, *RPF1*–*RPF3* (Bhattarai et al., 2022; Gao et al., 2022; She et al., 2018), *RPF11* (Dijkstra and Raedts, 2019), and *RPF12* (Dijkstra and Raedts, 2019), which confer resistance against at least 17 downy mildew of the 19 races of this pathogen reported in spinach. A neighbor-joining tree using SVs within this introgression region relationship in the 94 *Spinacia* accessions suggested that the DM loci originated from both wild species as donors (Fig. 5d), mostly from *S. turkestanica*, but 14 from *S. tetrandra* (Table S25).

We identified 9,047 insertions and deletions that are specific to the Sp_YY_v1 assembly, and then compared their frequencies in the 59 *S. oleracea*, 22 *S. turkestanica* and 13 *S. tetrandra* wild spinach accessions. A total of 338 and 5,921 SVs, respectively, had significantly higher or lower frequencies in *S. oleracea* than in either *S. turkestanica* or *S. tetrandra*, using Fisher’s exact tests. Using a FDR value < 0.05, this reduced to 282 candidate domestication SVs, which we term “domSVs”, in 245 genes (Fig. 5e and Table S26). These domSVs were significantly (Fisher’s exact text, *P* < 5.46×10^-10^) in the same terminal gene-rich chromosomal regions in which SVs were mostly found (Fig. S17). Similar pattern was also detected based on SNP-based selective sweep using two approaches (Fig. 5e).

These two SVs were associated with domesticated morphotypes. First, a 620 bp insertion (SV id: INS8694) downstream of the *SpCUC3* (*Te17S22m088490* within pericentromeric region of chromosome 2) gene is enriched in *S. oleracea* accessions (FDR= 7.41×10^-10^, Fisher’s exact test) (Fig. 5f, g, Table S26), and associated with decreased expression (Fig. 5h, Table S28). Silencing of *CUC3* in *Cardamine hirsuta* can change leaf margins from serrated to smooth (Blein et al., 2008), and the three *Spinacia* species have expression differences consistent with such an effect on their leaf type differences (Fig. 5i). A 204-bp deletion (SV id: DEL28738, Fig. 5j, Fig. S19, Table S26) was located in the CDS of the *SpWNK5* (*Te17S22m261760* within non-pericentromeric region of chromosome 6) gene (Fig. S20), whose *Arabidopsis thaliana* homolog (*AtWNK5*) reduces flowering time (Wang et al., 2008). Expression of *SpWNK5* at three different growth stages (see legend of Fig. 5k) in two plants with different flowering time suggest that this gene might be involved in the flowering process in spinach (Fig. 5k).

Previous studies have attempted to understand the genetics of spinach improvement during domestication by searching for SNPs showing associations with cultivated spinach morphotypes (Ma et al., 2022; Xu et al., 2017). Although many such SNPs have been detected, most such variants may not affect functions or have phenotypic effects Our analysis uncovered two examples of SVs (Fig. 5g–k), such variants are associated with the *SpCUC3* and *SpWNK5* that might be involved in determining leaf margin type and flowering time in spinach.

We find that LD is higher in the pericentromeric regions of *Spinacia* genomes, compared with other genome regions, and diversity is lower, indicating that genes in this region do not evolve independently (Fig. 3). Although gene densities in these regions are low in *Spinacia* species, they are clearly not too low to cause the predicted effects on diversity. However, LD decays within < 100 kb on most *S. tetrandra* chromosomes, even for the pericentromeric regions (and usually within about half this distance for the non-pericentromeric regions, see Fig. 3c). This suggests that recombination occurs occasionally in the former, and our recombination rate estimates support this conclusion (Fig. 3d). If so, pericentromeric regions cannot be assumed to be fully recombinationally inactive. It may therefore be possible to detect genes affecting domestication phenotypes, or at least regions including them, even if they are within such regions. If the regions do not include too many genes, the number of candidates for a given phenotype might not be too large for testing. Our study therefore illustrates the value of analyses using re-sequencing of multiple individuals, which has revealed important details of genome behaviour that would not be apparent from genome sequences of just one or two individuals.

## Acknowledgments

This work was supported by the Chinese Academy of Agricultural Sciences Innovation Project (CAAS-ASTIP-IVFCAAS, CAAS-ZDRW202103), and China Agricultural Research System (CARS-23-A-17).

## Author contributions

W.Q. designed the study. H.S., Z.L., D.C., and F.C., analyzed the data. H.S. wrote the manuscript. W.Q., Z.L., H.Z., Z.X. prepared the samples. W.Q., D.C., J.W., X.W., F.C., revised the manuscript. D.C. reformulated the manuscript.

## References

Arbel-Eden, A. and Simchen, G. (2019) Elevated Mutagenicity in Meiosis and Its Mechanism. Bioessays 41, e1800235.

Becher, H., Jackson, B.C. and Charlesworth, B. (2020) Patterns of genetic variability in genomic regions with low rates of recombination. Current Biology 30, 94–100. e103.

Begun, D.J. and Aquadro, C.F.J.N. (1992) Levels of naturally occurring DNA polymorphism correlate with recombination rates in D. melanogaster. Nature 356, 519–520.

Bhattarai, G., Olaoye, D., Mou, B.Q., Correll, J.C. and Shi, A.N. (2022) Mapping and selection of downy mildew resistance in spinach cv. whale by low coverage whole genome sequencing. Frontiers in Plant Science 13, 1012923.

Blein, T., Pulido, A., Vialette-Guiraud, A., Nikovics, K., Morin, H., Hay, A., Johansen, I.E., et al. (2008) A Conserved molecular framework for compound leaf development. Science 322, 1835–1839.

Bussell, J.J., Pearson, N.M., Kanda, R., Filatov, D.A. and Lahn, B.T. (2006) Human polymorphism and human-chimpanzee divergence in pseudoautosomal region correlate with local recombination rate. Gene 368, 94–100.

Chan, A.H., Jenkins, P.A. and Song, Y.S. (2012) Genome-wide fine-scale recombination rate variation in drosophila melanogaster. Plos Genet 8, e1003090.

Charlesworth, Sniegowski and Stephan (1994) The evolutionary dynamics of repetitive DNA in eukaryotes. Nature 371, 215–220.

Charlesworth, B., Morgan, M. and Charlesworth, D.J.G. (1993) The effect of deleterious mutations on neutral molecular variation. Genetics 134, 1289–1303.

Chen, Y.Y., Schreiber, M., Bayer, M., Dawson, l.K., Hedley, P.E., Lei, L., Akhunova, A., et al. (2022) The evolutionary patterns of barley pericentromeric chromosome regions, as shaped by linkage disequilibrium and domestication. the plant journal 111, 1580–1594.

Dijkstra, J.A. and Raedts, R. (2019) Spinach plants that are resistant to downy mildew.

Dolgin, E.S. and Charlesworth, B. (2008) The effects of recombination rate on the distribution and abundance of transposable elements. Genetics 178, 2169–2177.

Filatov, D.A. and Gerrard, D.T.J.G. (2003) High mutation rates in human and ape pseudoautosomal genes. Gene 317, 67–77.

Gao, G., Lu, T., She, H., Xu, Z., Zhang, H., Liu, Z. and Qian, W. (2022) Fine mapping and identification of a candidate gene of downy mildew resistance, RPF2, in spinach (Spinacia oleracea L.). Int J Mol Sci 23, 14872.

Hamblin, M.T., Casa, A.M., Sun, H., Murray, S.C., Paterson, A.H., Aquadro, C.F. and Kresovich, S. (2006) Challenges of detecting directional selection after a bottleneck: Lessons from Sorghum bicolor. Genetics 173, 953–964.

Hellmann, I., Ebersberger, I., Ptak, S.E., Pääbo, S. and Przeworski, M. (2003) A neutral explanation for the correlation of diversity with recombination rates in humans. Am J Hum Genet 72, 1527–1535.

Hickey, L.T., Hafeez, A.N., Robinson, H., Jackson, S.A., Leal-Bertioli, S.C.M., Tester, M., Gao, C.X., et al. (2019) Breeding crops to feed 10 billion. Nat Biotechnol 37, 744–754.

Jackson, B.C., Campos, J.L., Haddrill, P.R., Charlesworth, B. and Zeng, K. (2017) Variation in the intensity of selection on codon bias over time causes contrasting patterns of base compositionevolution in Drosophila. Genome Biology and Evolution 9, 102–123.

Jaillon, O., Aury, J.M., Noel, B., Policriti, A., Clepet, C., Casagrande, A., Choisne, N., et al. (2007) The grapevine genome sequence suggests ancestral hexaploidization in major angiosperm phyla. Nature 449, 463–U465.

Jayakodi, M., Padmarasu, S., Haberer, G., Bonthala, V.S., Gundlach, H., Monat, C., Lux, T., et al. (2020) The barley pan-genome reveals the hidden legacy of mutation breeding. Nature 588, 284–289.

Langley, S.A., Karpen, G.H. and Langley, C.H. (2014) Nucleosomes Shape DNA Polymorphism and Divergence. Plos Genet 10, e1004457.

Lin, T., Zhu, G., Zhang, J., Xu, X., Yu, Q., Zheng, Z., Zhang, Z., et al. (2014) Genomic analyses provide insights into the history of tomato breeding %J Nature Genetics. Nature Genetics 46, 1220–1226.

Ma, X.K., Yu, L.A., Fatima, M., Wadlington, W.H., Hulse-Kemp, A.M., Zhang, X.T., Zhang, S.C., et al. (2022) The spinach YY genome reveals sex chromosome evolution, domestication, and introgression history of the species. Genome Biology 23, 23–75.

Marais, G. and Galtier, N. (2003) Sex chromosomes: how X-Y recombination stops. Current Biology 13, R641–R643.

Mascher, M., Wicker, T., Jenkins, J., Plott, C., Lux, T., Koh, C.S., Ens, J., et al. (2021) Long-read sequence assembly: a technical evaluation in barley. Plant Cell 33, 1888–1906.

McVean, G., Awadalla, P. and Fearnhead, P. (2002) A coalescent-based method for detecting and estimating recombination from gene sequences. Genetics 160, 1231–1241.

Nielsen, R., Williamson, S., Kim, Y., Hubisz, M.J., Clark, A.G. and Bustamante, C. (2005) Genomic scans for selective sweeps using SNP data. Genome Res 15, 1566–1575.

Ou, S. and Jiang, N. (2018) LTR_retriever: a highly accurate and sensitive program for identification of long terminal repeat retrotransposons. Plant Physiology 176, 1410–1422.

Purugganan, M.D. (2019) Evolutionary insights into the nature of plant domestication. Current Biology 29, R705–R714.

Ribera, A., Treuren, R.V., Kik, C., Bai, Y., Wolters, A.J.G.R. and Evolution, C. (2020) On the origin and dispersal of cultivated spinach (Spinacia oleracea L.). Genetic Resources Crop Evolution 68, 1023–1032.

Roach, M.J., Schmidt, S.A. and Borneman, A.R. (2018) Purge Haplotigs: allelic contig reassignment for third-gen diploid genome assemblies. BMC Bioinformatics 19, 460.

Ross-Ibarra, J., Morrell, P.L. and Gaut, B.S. (2007) Plant domestication, a unique opportunity to identify the genetic basis of adaptation. P Natl Acad Sci USA 104, 8641–8648.

She, H., Liu, Z., Li, S., Xu, Z., Zhang, H., Cheng, F., Wu, J., et al. (2023) Evolution of the spinach sex-linked region within a rarely recombining pericentro-meric region. Plant Physiol 00, 1–18.

She, H.B., Liu, Z.Y., Xu, Z.S., Zhang, H.L., Cheng, F., Wu, J., Wang, X.W., et al. (2022) Comparative chloroplast genome analyses of cultivated spinach and two wild progenitors shed light on the phylogenetic relationships and variation. Scientific Reports 12, 856.

She, H.B., Qian, W., Zhang, H.L., Liu, Z.Y., Wang, X.W., Wu, J., Feng, C.D., et al. (2018) Fine mapping and candidate gene screening of the downy mildew resistance gene RPF1 in spinach. Theoretical and Applied Genetics 131, 2529–2541.

Stephan, W. (2019) Selective Sweeps. Genetics 211, 5–13.

Sun, P., Jiao, B., Yang, Y., Shan, L., Li, T., Li, X., Xi, Z., et al. (2022) WGDI: A user-friendly toolkit for evolutionary analyses of whole-genome duplications and ancestral karyotypes. Mol Plant 15, 1841–1851.

Sun, X., Jiao, C., Schwaninger, H., Caho, C., Ma, Y., Duan, N., Khan, A., et al. (2020) Phased diploid genome assemblies and pan-genomes provide insights into the genetic history of apple domestication. Nature genetics 52, 1423–1432.

Tajima, F. (1989) The effect of change in population size on DNA polymorphism. Genetics 123, 597–601.

Tian, J.G., Wang, C.L., Xia, J.L., Wu, L.S., Xu, G.H., Wu, W.H., Li, D., et al. (2019) Teosinte ligule allele narrows plant architecture and enhances high-density maize yields. Science 365, 658–664.

Treuren, R., Groot, L., Hisoriev, H., Khassanov, F., Farzaliyev, V., Melyan, G., Gabrielyan, I., et al. (2020) Acquisition and regeneration of Spinacia turkestanica Iljin and S. tetrandra Steven ex M. Bieb. to improve a spinach gene bank collection. Genetic Resources Crop Evolution 67, 549–559.

Wang, Y., Liu, K., Liao, H., Zhuang, C., Ma, H. and Yan, X. (2008) The plant WNK gene family and regulation of flowering time in Arabidopsis. Plant Biology 10, 548–562.

Waterhouse, R.M., Mathieu, S., A, S.F., Mosè, M., Panagiotis, I., Guennadi, K., Kriventseva, E.V., et al. (2017) BUSCO applications from quality assessments to gene prediction and phylogenomics. Molecular Biology and Evolution 3, 3.

Xu, C., Jiao, C., Sun, H., Cai, X., Wang, X., Ge, C., Zheng, Y., et al. (2017) Draft genome of spinach and transcriptome diversity of 120 Spinacia accessions. Nature Communications 8, 15275.

Zhang, W.Y., Tan, C., Hu, H.F., Pan, R., Xiao, Y.H., Ouyang, K., Zhou, G.F., et al. (2023) Genome architecture and diverged selection shaping pattern of genomic differentiation in wild barley. Plant Biotechnology Journal 21, 46–62.

