## Supplemental Figures for "Insights into spinach domestication from genomes sequences of two wild spinach progenitors, *Spinacia turkestanica* and *S. tetrandra*"

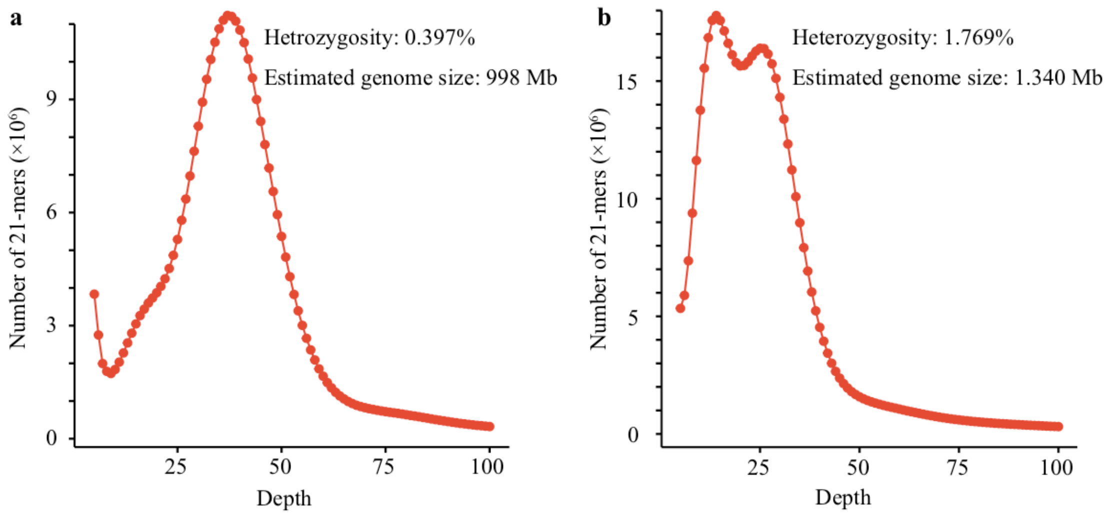


**Figure S1. 21-mer spectrum of Illumina reads of (a) *S. turkestanica* and (b) *S. tetrandra.***

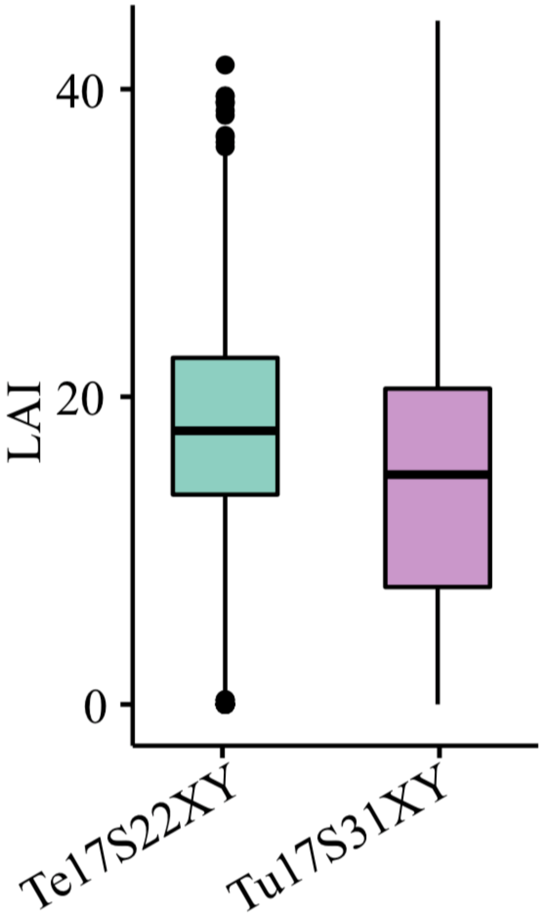


**Figure S2. LTR assembly index (LAI) of the two wild assemblies.**

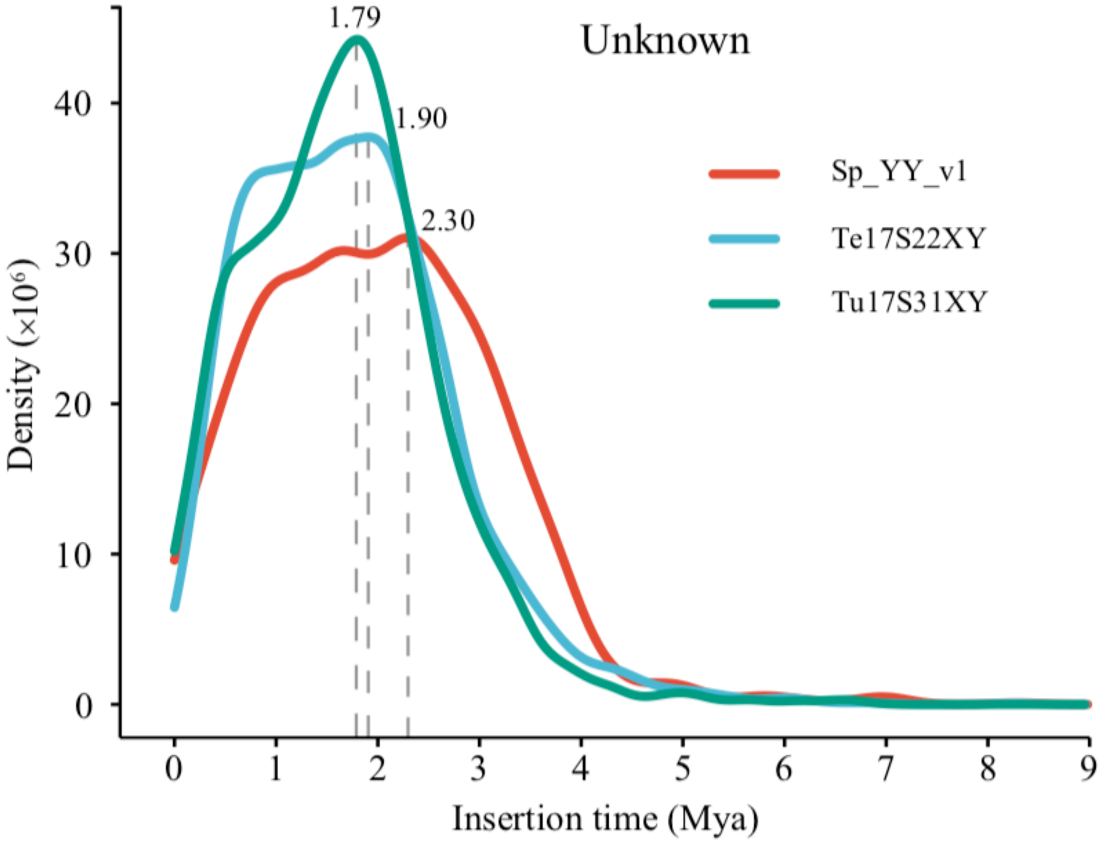


**Figure S3. Insertion time of unknown intact LTR-RT among the *S. tetrandra*, *S. turkestanica*, and *S. oleracea*.** Insertion time were inferred based on a mutation rate of $7.0 \times 10^{-9}$ substitutions per site per year. MYA: million years ago.

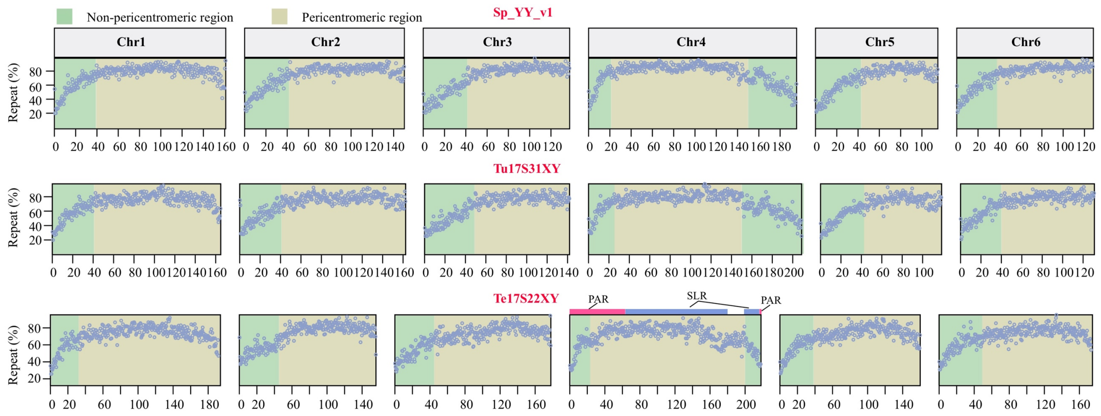


**Figure S4. Identification of pericentromeric regions for each chromosome using change-point analysis of repeat densities estimated in 500-kb windows**. PAR: pseudo-autosomal region; SLR: sex-linked region.

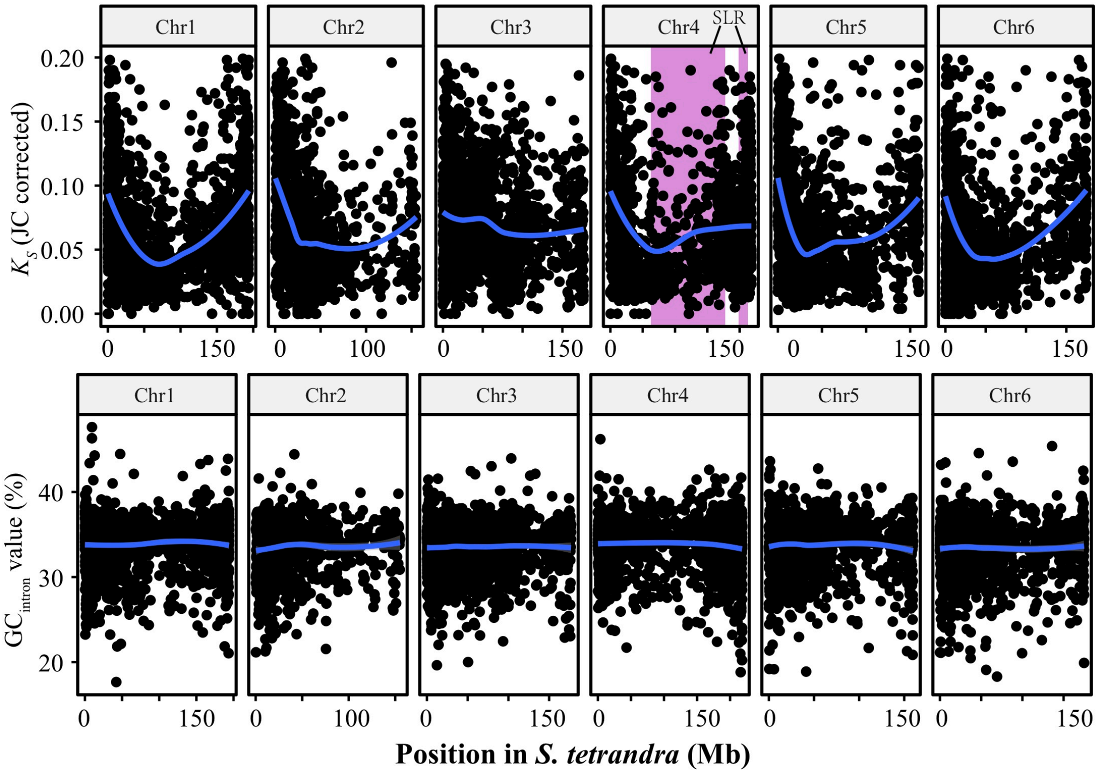


**Figure S5. *Ks* and GC values of single-copy genes in *S. tetrandra* and their orthologs in the Tu17S31XY assembly.** The GC values were computed as percentages, using the introns of genes in the *S. tetrandra* assembly. Single-copy genes with coding sequences > 200 bp and intron length > 100 bp were used for both the *Ks* and the GC analyses. The sex-linked regions of chromosome 4 in *S. tetrandra* are indicated in pink. These regions show a clear tendency to have lower inter-species divergence than the rest of chromosome 4.

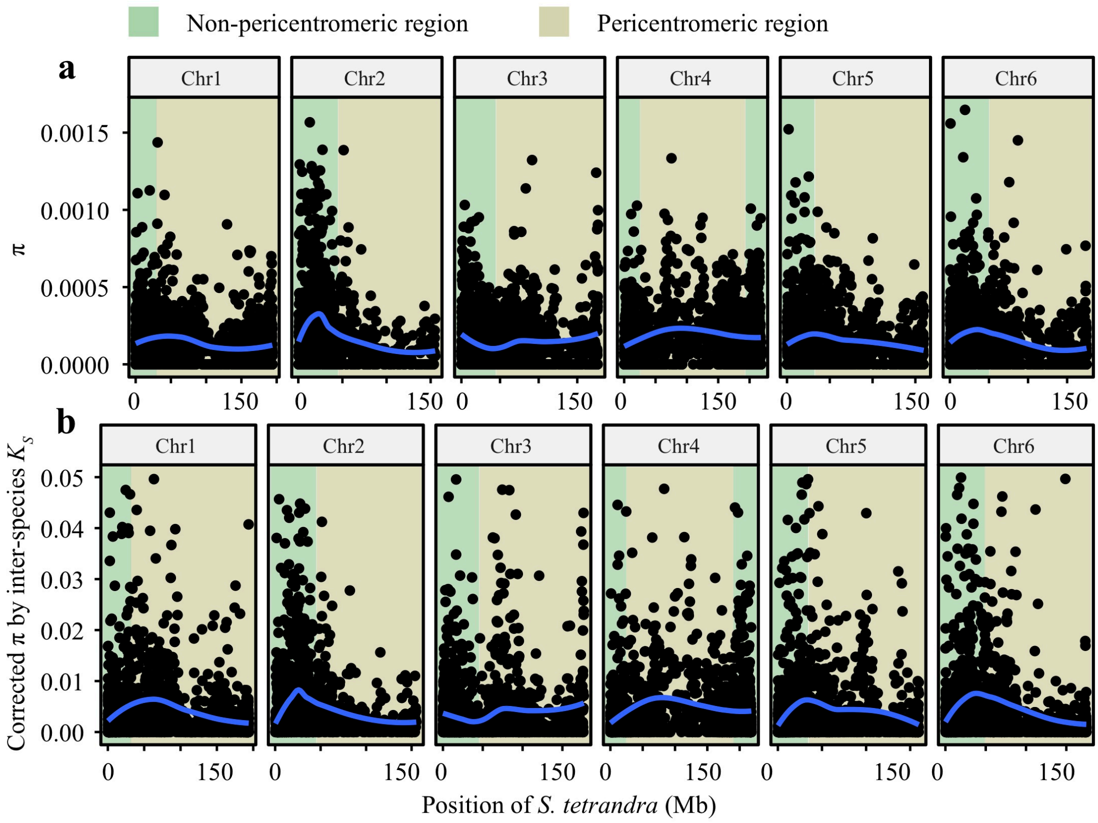


**Figure S6. Distribution of nucleotide diversity ($\pi_S$) values for synonymous sites for single-copy genes in 13 *S. tetrandra* accessions.** Part **(a)** shows the $\pi_S$ values, and **(b)** shows values corrected by the *S. tetrandra*-*S. turkestanica* synonymous divergence values for the same genes. Each dot represents the value for a single-copy gene. The smoothed lines were plotted using Loess method in R.

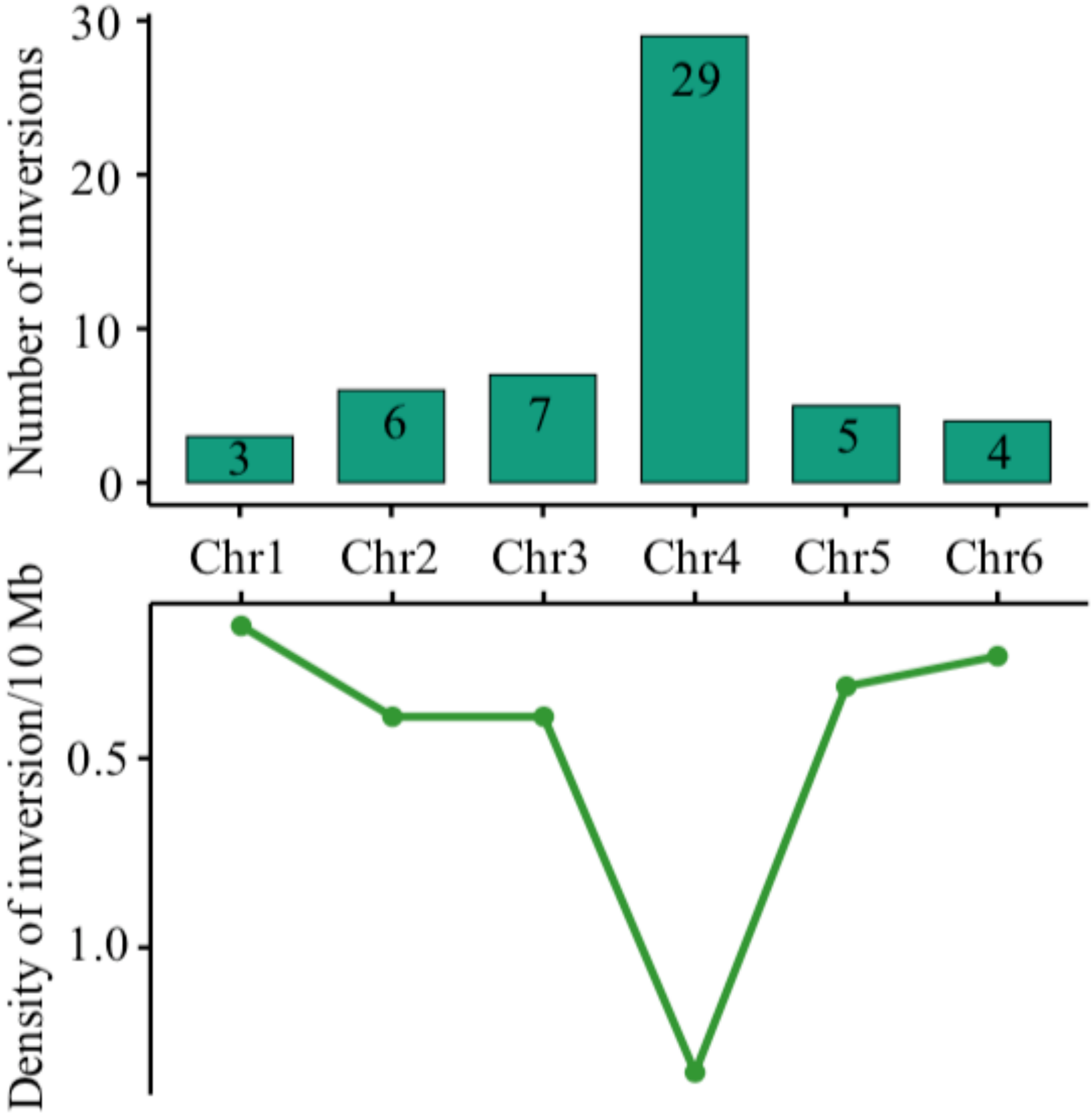


**Figure S7. Distribution and density of non-redundant large inversions, of >1 Mb, in the Te17S22XY assembly.**

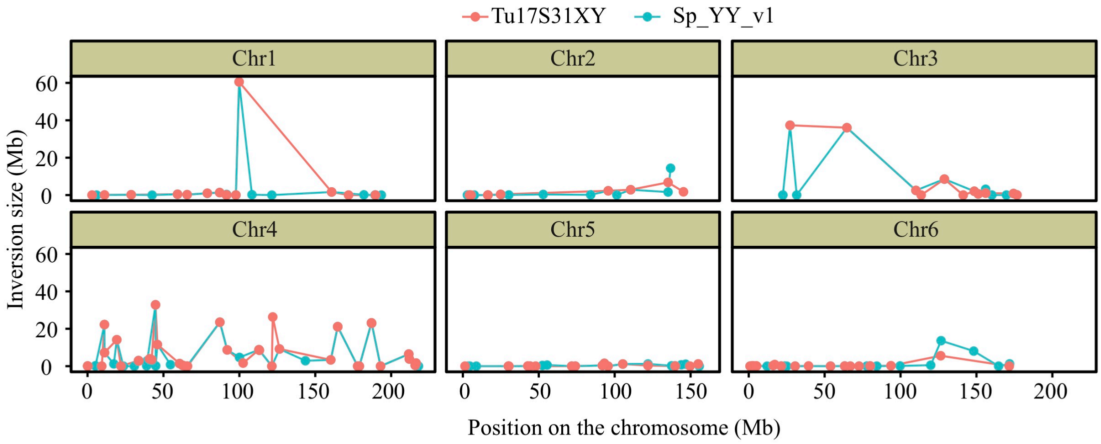


**Figure S8. Distribution of inversion sizes in the Te17S22XY assembly, relative to the two other *Spinacia* reference genome sequences indicated at the top of the figure.** Each dot indicates an inversion.

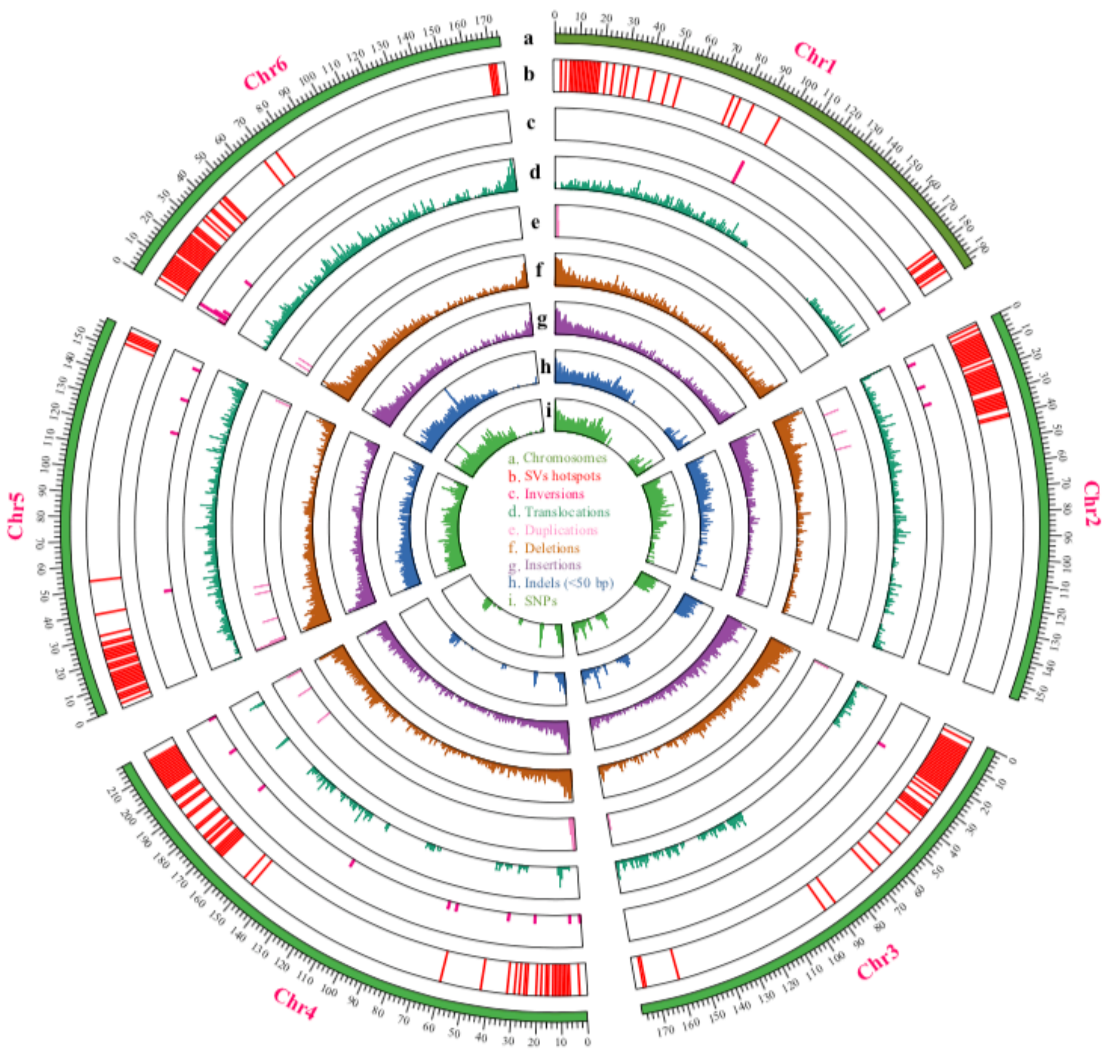


**Figure S9. Landscape of non-redundant variants on the Te17S22XY assembly. Densities of variants are estimated within 100-kb windows.**

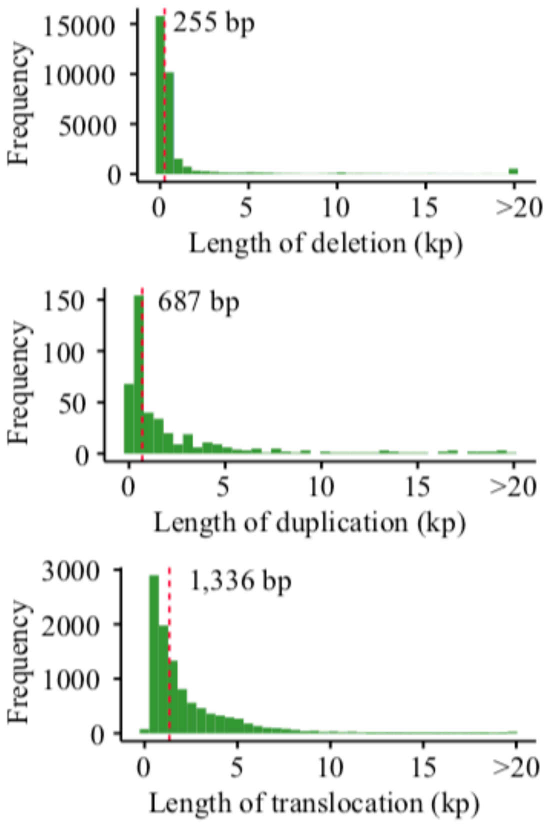


**Figure S10. The frequency of SVs (deletion, insertion, duplication, inversion, and translocation).** Dotted red lines represent the median lengths of the SVs.

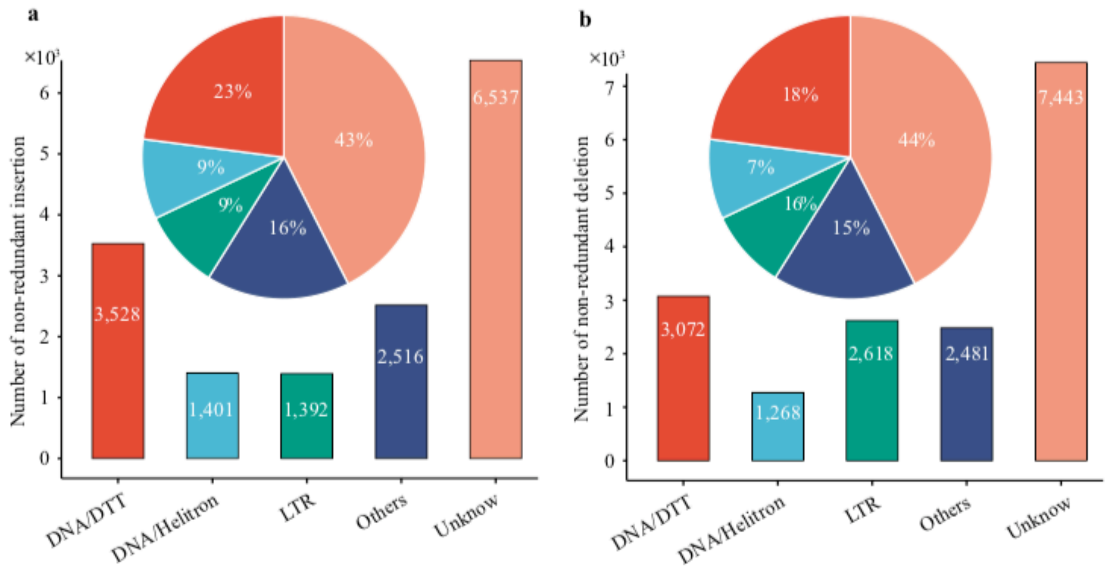


**Figure S11. The number and percentage of non-redundant (a) insertions and (b) deletions that related to TEs.**

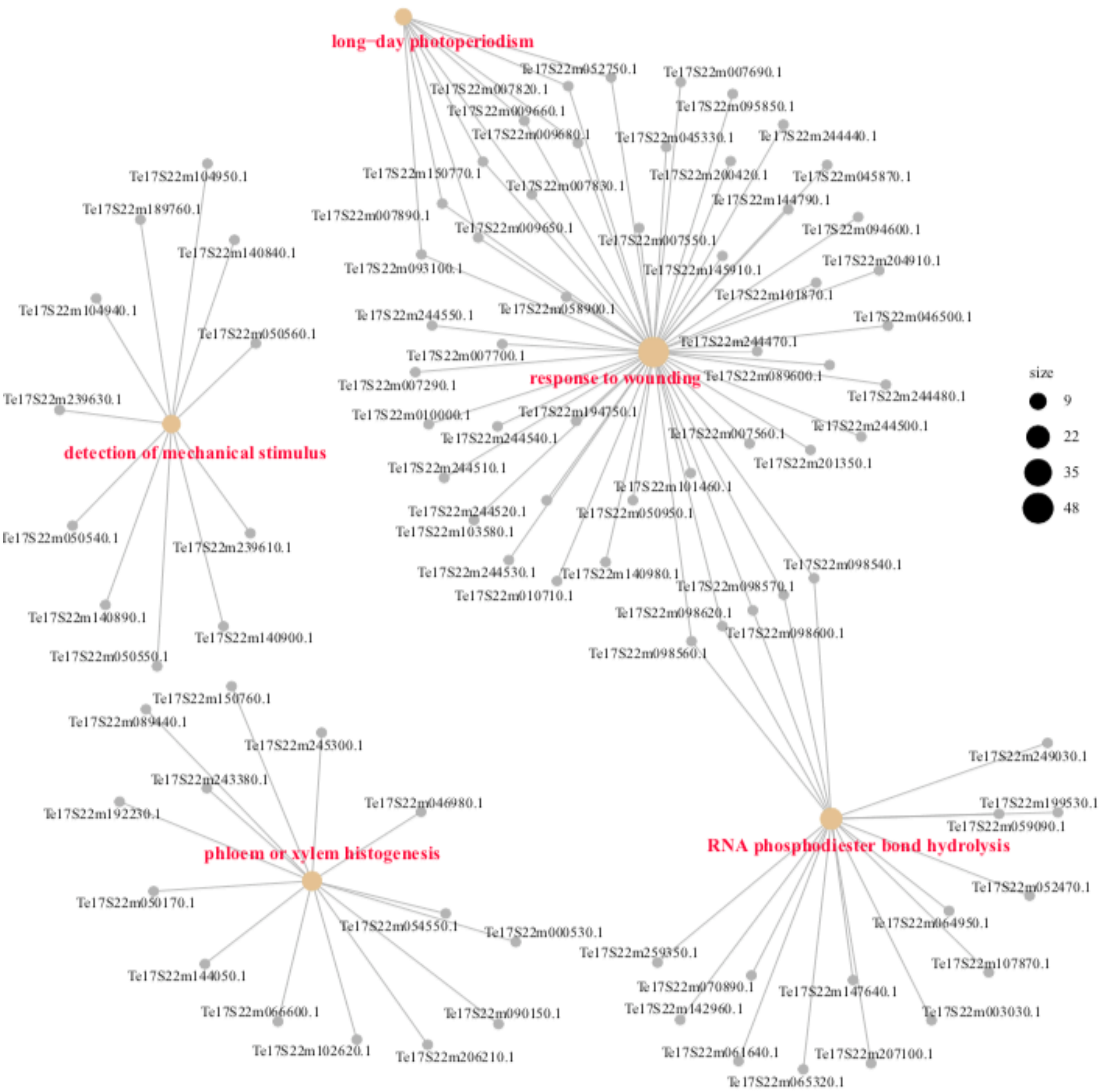


**Figure S12. Enrichment of gene ontology (GO) under the "biological process" category for the genes within hotspot regions. $p < 0.05$, calculated using Fisher's exact test.** Circle size represent the frequency of each enriched GO term.

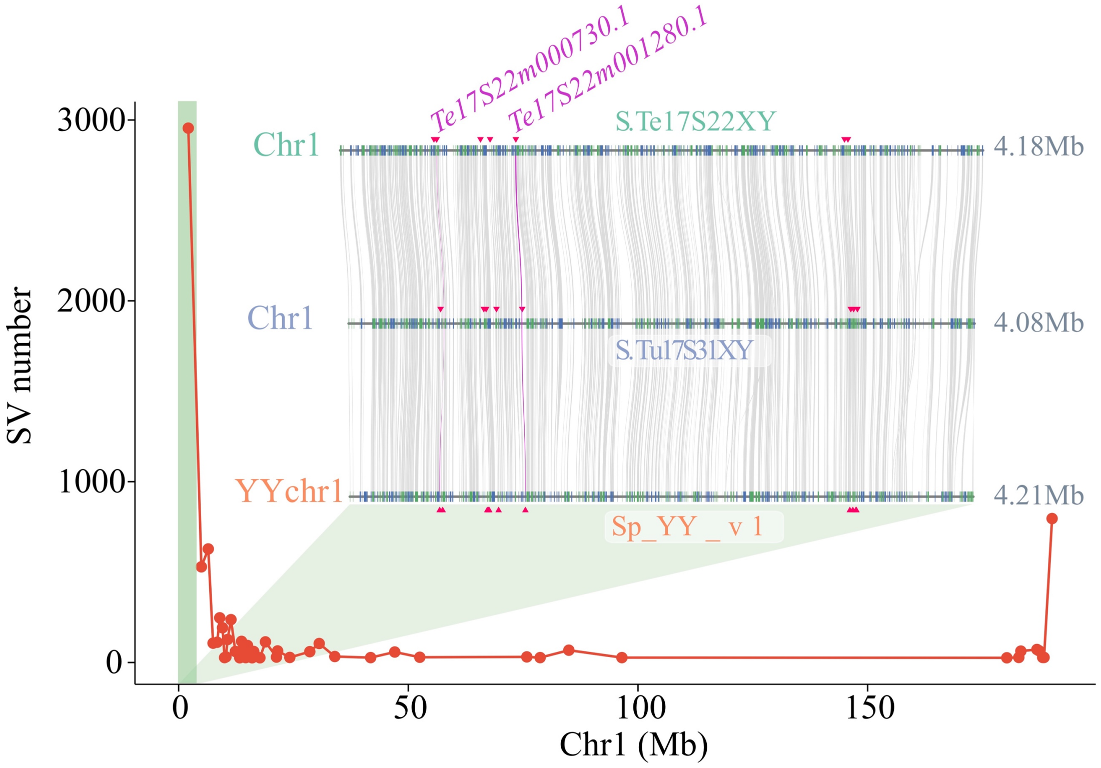


**Figure S13. Distribution of SV hotspot regions on the *S. tetrandra* chromosome 1.** Each dot represents a hotspot region. A representative hotspot region, from 0–4.2 Mb, is highlighted in light green and shown in detail in the three *Spinacia* species assemblies were used for microsynteny analysis. The blue and green vertical lines on the chromosome diagrams indicate genes on the plus and min strand, respectively. The gene with an NBS/NBS-LRR domain is marked with red triangles. Three reported spinach candidate genes, for downy mildew resistance, *RPF1* (*Te17S2m000730*), *RPF2* and *RPF3* (*Te17S22m001280*) (*RPF2* and *RPF3* have the same candidate genes as reported previously, suggesting that the two genes are close) are highlighted in purple.

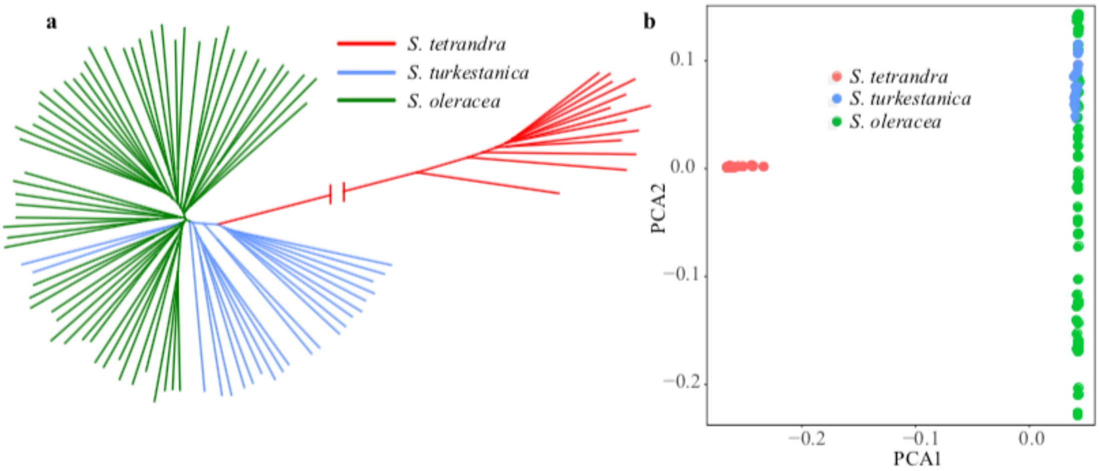


**Figure S14. Phylogeny and PCA plot of cultivated spinach and its wild relatives.** Phylogenetic tree **(a)** and PCA plot **(b)** of 13 *S. tetrandra*, 22 *S. turkestanica*, and 59 *S. oleracea* accessions based on 982,999 SNPs.

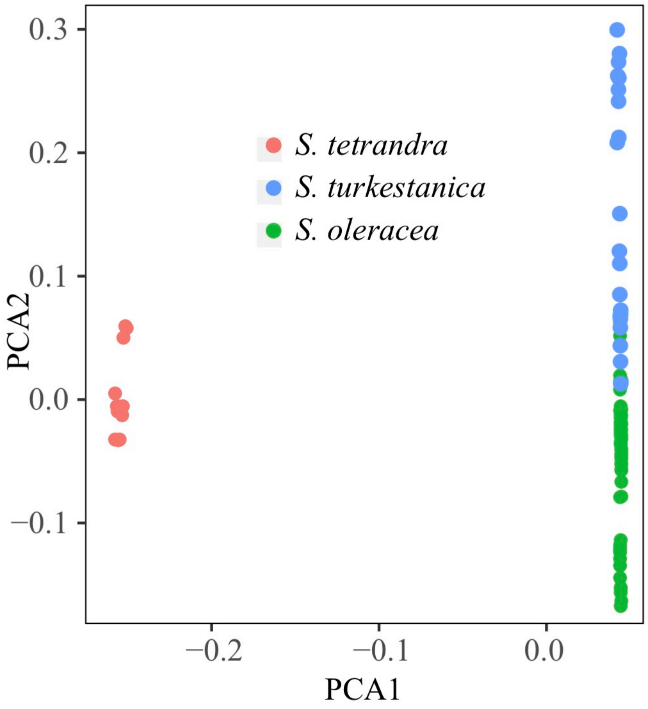


**Figure S15. PCA plot of 13 *S. tetrandra*, 22 *S. turkestanica*, and 59 *S. oleracea* accessions based on SVs (Insertion and deletion).**

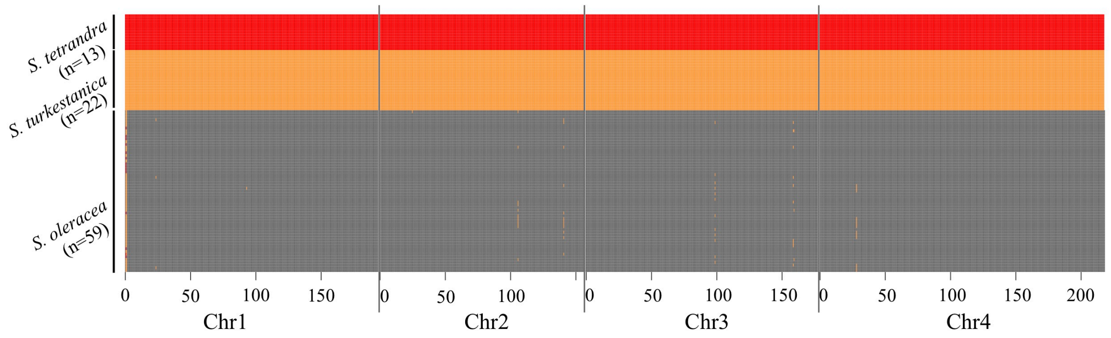


**Figure S16. Identification of introgressed region from the two wild relatives of cultivated spinach.** Species-specific genome regions are shown in different colors. Note that chromosomes 5 and 6 are not shown because we do not detect introgression region on these chromosomes.

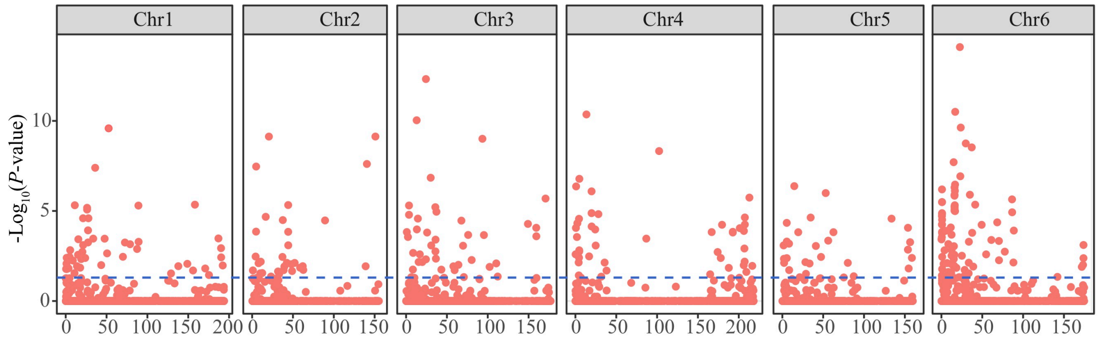


**Figure S17. Potentially selected SVs in *S. oleracea*, based on their frequencies compared with those in the two wild relatives.** We ascertained insertions and deletions in the cultivated spinach genome assembly, and then calculated their frequencies in the cultivated spinach samples, and in our samples of its two wild relative accessions, and performed Fisher's exact tests for each SV, to detect ones whose frequencies are different in cultivated spinach. The blue dotted line indicates the FDR value < 0.05.

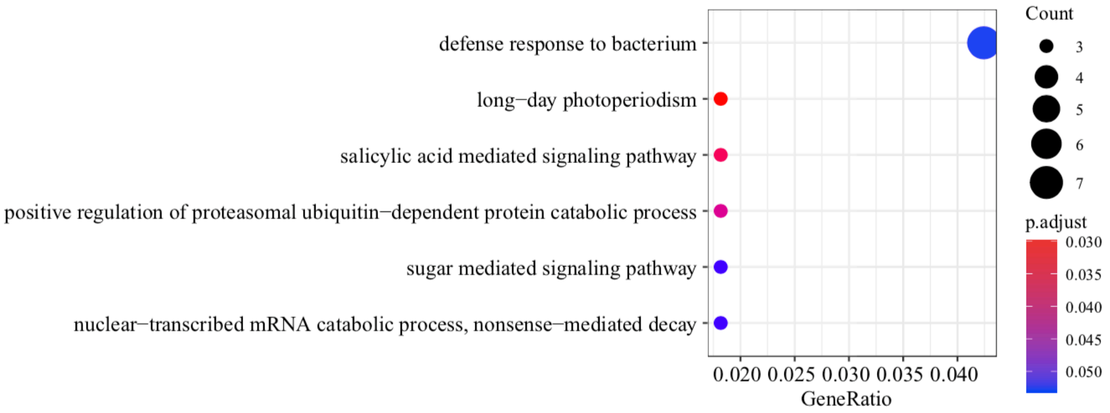


**Figure S18. Enrichment of gene ontology (GO) under the "biological process" category for the genes related to the domestication of *S. oleracea*. $p < 0.05$, calculated using Fisher's exact test.** Circle size represent the frequency of each enriched GO term.

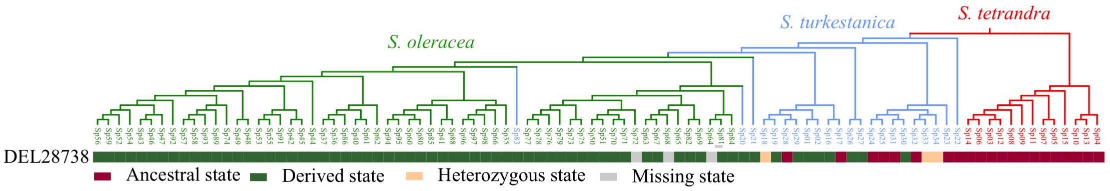


**Figure S19. The distribution of DEL28738 genotype in 94 *Spinacia* accessions.** For an SV state that is the same as *S. tetrandra* is defined as the ancestral state, otherwise that is referred as derived state.

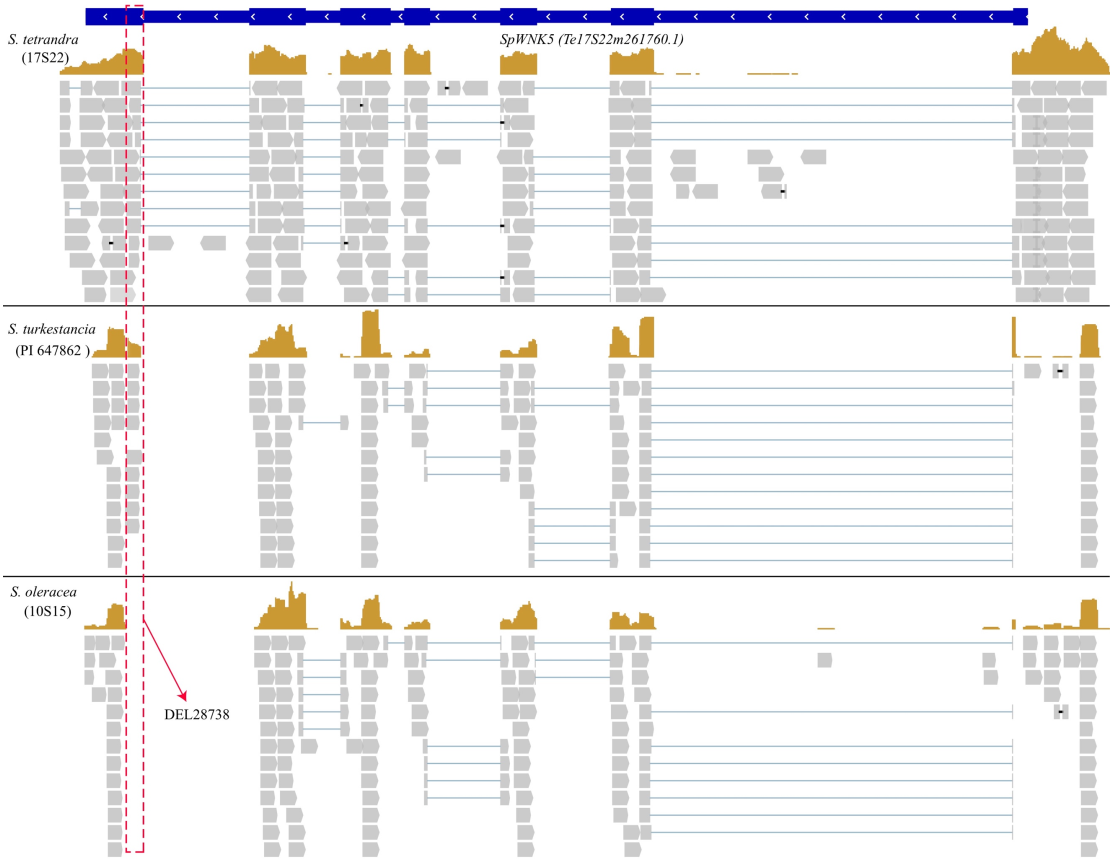


**Figure S20. Validation of DEL28738 using RNA-seq reads from *S. tetrandra*, *S. turkestanca* and *S. oleracea* accessions.** The RNA-seq data of *S. turkestanica* is collected from Xu et al., (2017).
